# Objective method to estimate the duration of the DNA replication cycle without perturbation in *Escherichia coli*

**DOI:** 10.64898/2026.02.10.705063

**Authors:** Alix Meunier, Florian Sarron, Isabelle Bru, Meriem Benkhelifa, Erick Denamur, Manuel Campos

**Affiliations:** Laboratoire de Microbiologie et de Genetique Moleculaires, Centre de Biologie Integrative, Universite de Toulouse, CNRS, Toulouse, France; IRIT, CBI, CNRS, Université de Toulouse, Toulouse, France; Universite Paris Cite, Universite Sorbonne Paris Nord, INSERM, IAME, Paris, France

## Abstract

All cells, including unicellular organisms such as *Escherichia coli*, depend on the robustness of their cell cycle for proliferation. The cell cycle is central to the organism’s highly resilient proliferation program and is thus a key focus in many areas of cell biology. However, studying the DNA replication cycle in bacteria remains challenging due to technical limitations. The durations of the replicative and post-replicative phases (known as the C and D periods, respectively) are inferred from the analysis of cellular DNA content distributions, obtained through microscopy or flow cytometry. This analysis typically requires pharmacological treatments to conduct replication run-off experiments, which help distinguish between cells that have or have not yet initiated DNA replication.

Four decades ago, Skarstad, Steen and Boye showed that flow cytometry profiles measuring DNA amount per cell from exponentially growing cells could provide sufficient information to infer the durations of cell cycle periods at low growth rates. In this study, we propose an objective and automated approach that implements this idea to estimate cell cycle parameters for any growth conditions. Specifically, we validate a Nested Sampling method to estimate cell cycle parameters directly from flow cytometry data, eliminating the dependence on bacterial strain sensitivity to drugs. This tool, available as a [Python package](https://src.koda.cnrs.fr/genome-dynamics/bacycle), allows for the accurate and minimally biased estimation of the C+D period under any biologically relevant conditions. Given its independence from pharmacological treatments, we anticipate broad adoption of this tool, especially as we show that most natural isolates of *E. coli* are not amenable to the state of the art replication run-off experiments.

## Introduction

Bacteria evolved over 3.5 billion years to become the highly integrated and very robust proliferative agents we encounter today everywhere on earth. Some of them evolved as specific pathogens for humans and plants while most of them participate to ecosystems’ dynamics. Studying their proliferation process is a cornerstone of microbiology, with wide-reaching implications across various scientific disciplines and practical applications, making it a critical area of research. The robustness of cell proliferation is determined by a tightly controlled cell cycle progression. The bacterial cell cycle is defined as a succession of periods, known as the B, C, and D periods, similar in essence, but mechanistically different from the G1, S and G2/M phases of the eukaryotic cell cycle (see note in [1]). The interval between the completion of cell division and the initiation of DNA replication defines the B period. The C period is the time period necessary for the duplication of the bacterial chromosome. This is a highly regulated and precise process ensuring that the genetic material is accurately copied [2–4]. Finally, the D period encompasses the time between the end of DNA replication and the division of the cell into two daughter cells. The ability of some bacterial species to initiate new rounds of DNA replication before cell division, or even before the end of the ongoing replication period allows for a generation time *τ* (time between 2 successive divisions) shorter than the time required to replicate and segregate DNA (*C* + *D > τ*).

In the past, these successive periods have been defined using radiolabeled thymine pulse-chase experiments coupled to a membrane elution technique [5] or elutriation [6] to measure the rate of radiolabeled thymine incorporation at any given cell age [7, 8]. Kubitschek and Newman estimated the C and D periods in slow growing *E. coli* without any cell sorting method. Instead, they used the DNA damage related lethal effect of the radio isotope *I*^125^ when incorporated during replication in bulk population [9]. Flow cytometry was later used to quantify DNA amount per cell in asynchronous cell populations.

To this date, the state-of-the-art approach to estimate these periods remains to use a combination of rifampicin and cephalexin to perform replication run-off experiments coupled to flow cytometry. The typical replication run-off experiment is based on the ability of rifampicin to inhibit new rounds of initiation of DNA replication while allowing the ongoing ones to terminate [10]. This type of experiment has been widely used on laboratory strains over decades using rifampicin or chloramphenicol [11]. Rifampicin primarily inhibits transcription. However, probably as a combination of effects on DnaA protein production [12] and DNA topology in the vicinity of the origin of replication [13], rifampicin also prevents any new initiation of DNA replication, while allowing ongoing replication cycles to terminate. As a result of rifampicin addition, an asynchronously growing population of cells split up in two sub-populations: the cells that didn’t start their DNA replication at the moment of antibiotic treatment end up with *n* chromosome(s), and the cells that did initiate replication and end up with 2*n* chromosomes. Cephalexin inhibits cell division and prevents the DNA content per cell from being halved. These 2 sub-populations appear as characteristic peaks on a flow cytometry profile. The proportion of cells with 2*n* (or *n*) chromosomes corresponds to the proportion of cells that passed (or did not passed) the replication initiation event at the time of treatment. Assuming an exponentially growing cell population at steady state, this proportion is the only parameter, with *τ*, necessary to calculate the duration of the C+D period relative to the generation time according to equations:

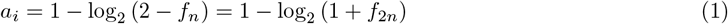

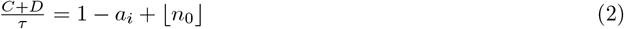

where *a*_*i*_ is the age at the time of initiation of DNA replication,*f*_*n*_ and *f*_2*n*_ are the fraction of cells with *n* or 2*n* chromosome equivalent(s) and ⌊*n*_0_⌋ an integer value equal to the number of overlapping replication cycles. Alternatively, the stringent response has been known to inhibit the initiation of new rounds of DNA replication for decades [14–16], probably through indirect effects on DnaA production as well as *dnaA* down-regulation [12]. Serine hydroxamate mimics a shortage in amino acids and triggers the stringent response, and induces replication run-off as a result [17]. However, the replication run-off effect induced by the serine hydroxamate treatment appeared to be less stringent than rifampicin treatment [14].

As early as 1985, an algorithm was proposed to compare the experimental cytometry profiles to theoretical profiles derived from the cell cycle model proposed in 1968 [7, 18]. The studies by Skarstad *et al*., (1985) Michelsen *et al*., (2003) and Stokke *et al*., (2012) [18–20] show that DNA distributions from flow cytometry on untreated cells contain enough information to accurately estimate cell cycle periods, at least for slow-growing *E. coli* cells. Yet, this approach remains poorly applied to studying the bacterial cell cycle, even for *E. coli*, likely because it requires a manual exploration of the parameter space.

Under slow growth conditions, the proportions of cells in B and D periods (*i*.*e*., non-replicating cells) result in very characteristic distributions of cellular DNA contents. Under richer growth conditions, the characteristic structures of the distribution become more subtle [19, 20]. In fact, no automated objective optimization method has been applied to estimate parameter values. One probable reason is that at its core the cell cycle model proposed by Cooper & Helmstetter is a piecewise model, and as such is not readily treated with standard optimization algorithms (typically using differentiation under the hood). Nevertheless, other methods are available to overcome the limitations associated with this challenge. Monte Carlo approaches are well-suited to optimize such problems, including Markov Chain Monte Carlo (MCMC) methods, if a unique solution exists, or more recently, Nested Sampling [21, 22] that has the advantage to take into account potential multiple solutions in case different sets of parameters would result in very similar outputs (*e*.*g*., multiple sets of cell cycle period values resulting in highly similar flow cytometry profiles).

More generally, Nested Sampling is a powerful Bayesian inference algorithm that can be used to estimate parameter posteriors and marginal likelihoods for complex models given observed data. In brief, a Bayesian algorithm starts with a guess about the parameter values (the prior belief), then update this *a priori* knowledge by gathering evidence (experimental data), and using a systematic method (here, the Nested Sampling algorithm) to update the belief toward parameters that best describe the data under the model. Remarkably, the Nested Sampling algorithm is able to efficiently capture multiple sets of parameters as solutions if they describe equally well the data (*i*.*e*., it is possible to efficiently sample multimodal posterior probabilities) [23]. This property can be used to test whether the classical cell cycle model is identifiable or not, and to refine the parameter search in constrained parameter range values, if necessary. In other words, it can be used to verify the unicity of any solution, or alternatively define a region in the parameter space where a unique set of parameters can describe the experimental data with the model.

We propose here to complete past efforts by providing the community with a tool based on the standard Cooper-Helmstetter DNA replication model coupled to a fitting procedure based on Nested Sampling [21, 22]. An alternative MCMC approach is also implemented in *bacycle*. In this study we first emphasize the need for such a tool by showing the limited spectrum of natural isolates amenable to classical replication run-off. We then propose and validate a Nested Sampling approach to estimate DNA replicative periods for bacteria and benchmark *bacycle* with replication run-off results.

## Materials and methods

### Strains and growth conditions

Bacteria were grown in a BSL2 laboratory, in 14mL culture tubes (ThermoScientific, USA) at 37^°^C with agitation, in the growth media indicated in the text. Bacterial strains and growth media used in this study are listed in Table 1 and 2, respectively. Strains were isolated from frozen stocks on LB-agar plates and a single colony was picked to inoculate 2mL of LB to grow an overnight pre-culture. The following morning, exponential precultures were inoculated from overnight precultures at a dilution of 1:500 and used to inoculate cultures when they reached an optical density at 600nm (OD_600nm_) close to 0.1. Cells were sampled when cultures reached an OD_600nm_ between 0.06 and 0.1. Rifampicin and cephalexin were added at 150*µ*g/mL and 10*µ*g/mL final from 30mg/mL and 10mg/mL stocks, respectively. Antibiotic treatments were conducted for 2 to 3 hours before sampling.

**Table 1.**
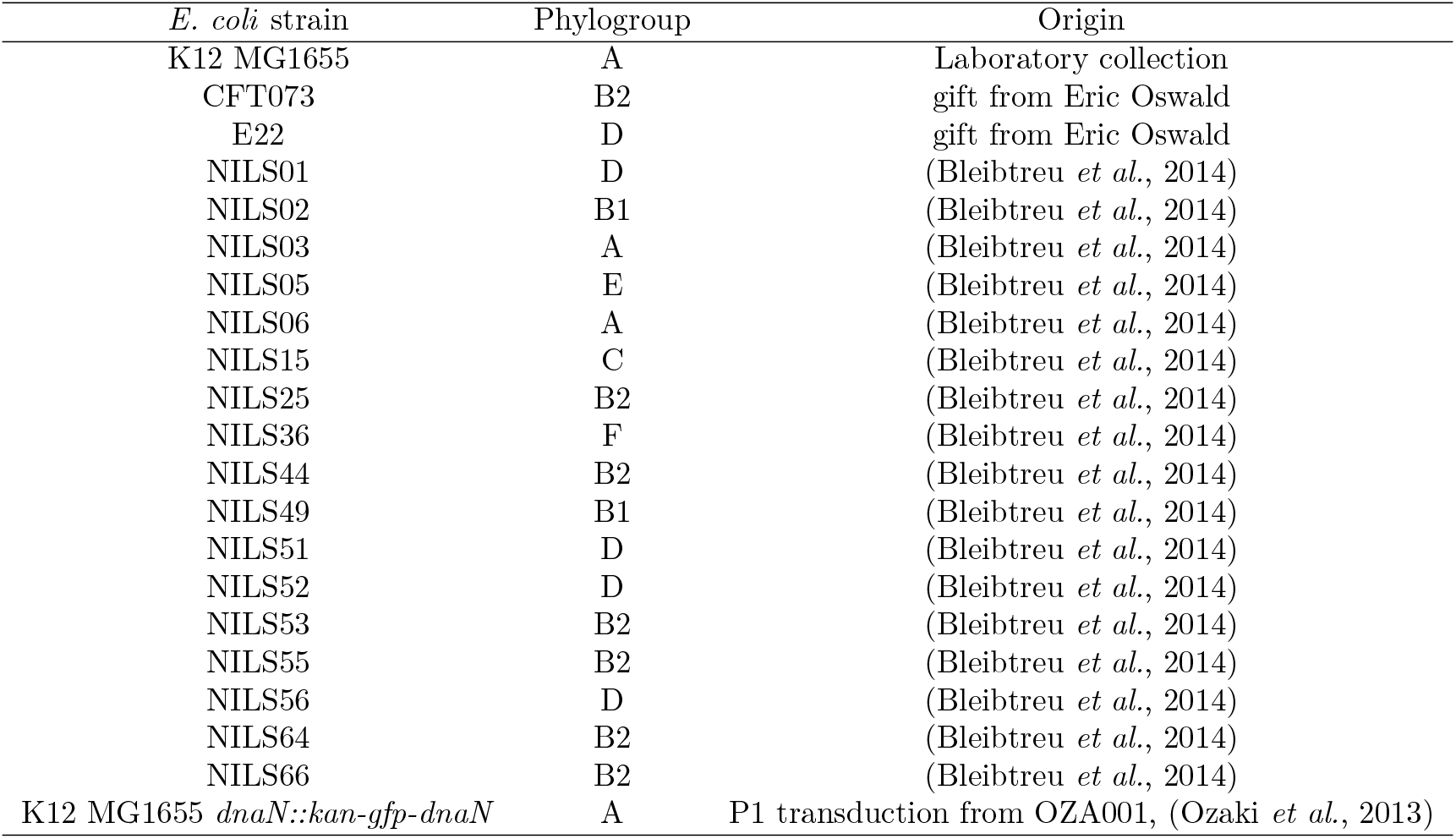
*E. coli* strains used in this study.

**Table 2.** Growth media used in this study and their composition.

| Medium | Composition |
| --- | --- |
| LB | Lysogeny Broth (Euromedex) |
| M9_Glu_CSA | M9 medium supplemented with glucose 0.2% w/v, casamino acids 0.1% w/v |
| M9_Ara_CSA | M9 medium supplemented with arabinose 0.2% w/v, casamino acids 0.1% w/v |
| M9_Glu_LB5% | M9 medium supplemented with glucose 0.2% w/v, LB 5% (v/v) |
| M9_Glu | M9 medium supplemented with glucose 0.2% w/v |
| M9_Ara | M9 medium supplemented with arabinose 0.2% w/v |
| M9_Gly_CSA | M9 medium supplemented with glycerol 0.2% w/v, casamino acids 0.1% w/v |
| M9_Gly | M9 medium supplemented with glycerol 0.2% w/v |

P1 transduction were conducted in accordance with conventional methods [24]. Briefly, a P1*vir* phage lysate was prepared on strain OZA001 [25] by mixing 300*µ*L of an overnight culture with 2*µ*L of P1*vir*. The resulting phase lysate was mixed at different multiplicities of infection with *E. coli* K12 MG1655 cells for 15min in LB at 37^°^C, and the infection was stopped by the addition of sodium citrate (7.5mM final) and cells were further incubated for 1h at 37^°^C before plating on selective plates (20*µ*g/mL kanamycin, 7.5mM sodium citrate). One clone from the smallest multiplicity of infection condition was further characterized by the formation of green fluorescent profiles under the microscope and by PCR.

### Growth curves

#### Pre-culture, dilutions and acquisition

Pre-cultures were grown to stationary phase in 96-well plates (first column of the plate), with each well inoculated from a single colony grown on LB agar. The pre-culture plates were incubated at 37^°^C, shaken at 180rpm for a minimum of 24 hours. The pre-cultures were then serially diluted in 5 wells per condition, typically to final dilution factors equal to 4000, 6000, 9000, 13500 and 20250. An exception was made for cultures in M9 Glycerol, which were inoculated from dilution factors from 1000, 2000, 3000, 4000 and 5000. The OD_600nm_ of these final dilutions was monitored over time using a FLUOstar Omega 96-well plate reader (BMG Labtech, France). A measurement of OD_600nm_ was taken every 10min, during 20h except for the cultures in M9 glycerol, which were carried out for 24h.

#### Growth curve analysis

The curves were smoothed by a LOESS function centered on 12 sliding points and the blank value is determined well by well (minimum value of the OD_600nm_ over the first 2 hours of acquisition, excluding the first measurement). The exponential part of each curve was determined by calculating the instantaneous growth rate (IGR, see formula below) and visually determining the plateau of IGR as a function of OD_600nm_.

The instantaneous growth rate (*IGR*) is calculated at each time point (*t*) according to the formula:

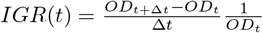

Where OD_t_ is the blank subtracted and smoothed optical density value at time point *t* and Δ*t* is the difference between two consecutive time points (here, 10min).

When *IGR*(*t*) was stable for at least 5 consecutive time points (50min), forming a plateau, the culture was considered in steady state and the logarithm of the OD_600nm_ values in this time window were used to fit with a linear regression model of the form:

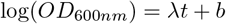

Where *λ* is the population growth rate and *b* the intercept (proxy for the log(OD_600nm_) of the inoculum). The population growth rate *λ* relates to the specific growth rate *µ* according to the formula *µ* = ln(2) · *λ*.

### Flow cytometry

#### Sample preparation

The cells were fixed in cold 70% ethanol for 1h and washed twice in filtered (pore size 0.22*µ*m), autoclaved PBS. The cells were then diluted to reach a final concentration below 6000 particles/*µ*L (typically a factor 2 between the culture and the sample analyzed by cytometry for exponential phase cultures, and a factor 500 for stationary phase cultures). The diluted cells are then incubated in staining buffer (10mM Tris-HCl, 10mM MgCl_2_, 100*µ*g/mL Mitramycin A (ref.M6891 Sigma-Aldrich, Merck, Germany) and 40*µ*g/mL Ethidium Bromide (ref.4905006 Dominique Dutscher, France) for 60min on ice.

#### Data acquisition

A sample of cells grown up to exponential phase in minimal medium M9 Glycerol was used as a standard in parallel replicates of each condition. Flow cytometry analysis was performed using a CytoFLEX instrument (Beckman Coulter Life Sciences, USA) in tube configuration. Samples were run at 10*µ*l/min. The primary detection threshold was set manually on the SSC-violet channel at 1500 arbitrary units. FSC and SSC signals were collected on the violet laser channel at 405nm with gains set to 98 and 101, respectively. DNA was detected by FRET. Mythramycin was excited at 405±nm and the Ethidium Bromide fluorescence signal was collected at 610±45nm with the gain of set to 2445. Particles corresponding to cells were gated according to their FSC and fluorescent signals (area) (see Figure S6 for gating strategy).

### Droplet digital PCR

#### DNA Extraction

DNA was isolated using the Wizard(r) Genomic DNA Purification Kit (Promega, USA) according to the instructions provided by the furnisher for Gram-negative bacteria, on culture sample in exponential growth phase. Extracted DNA is quantified with a NanoDrop (NanoDrop TM 1000 Spectrophotometer, ThermoFisher Scientific, USA) and extracts were diluted in distilled water to a final concentration of 1ng/*µ*L.

#### ddPCR

All experiments were performed using BioRad (USA) reagents and equipment QX200. A ddPCR reaction contains 11*µ*L of EvaGreen SuperMix QX200 ddPCR for primers, 152nM of each “ter” primer or 38nM of each “ori” primer, 0.25*µ*L of HindIII enzyme, 1*µ*L of template DNA at 1ng/*µ*L, and distilled water to the final volume of 22*µ*L per well. For *oriC* amplification, the following primers from [26] were used to amplify a 131bp region in teh *atpB* gene :

5’-GCCGCAGGATTACATAGGAC-3’ and 5’-CCACCGAGAAGAACATGGAG-3’

For *ter* amplification, the following primers from [27] were used to amplify a 175bp region in teh *ynfE* gene :

5’-AACTACGCGGGAAATACCC-3’ and 5’-TATCTTCCTGCTCAACGGTC-3’

The run consists of 34 cycles of denaturation (30sec at 95^°^C), primers annealing (30sec at 54^°^C) and extension (30sec at 60^°^C) with a 2^°^C/sec rate of temperature change. The raw data are ultimately analyzed with Quantasoft 1.7.4 software, allowing manual or automatic thresholding. The mean number of DNA targets per reaction were estimated with Poisson approximation according to the equation 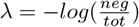, with *λ* the Poisson parameter (in copy/reaction) and *neg* and *tot* the numbers of negative and total partitions in the reaction (see Figure S7).

### Fluorescence microscopy

#### Image acquisition

Bacterial pre-cultures were inoculated from overnight cultures at a targeted density of 1-5 mOD_600nm_ to allow for 4 to 5 generations before use. Upon reaching OD_600nm_ within the 0.05-0.1 range, cultures were inoculated from the precultures at a targeted density of 15 mOD_600nm_ to allow for an extra 2 to 3 generations before sampling cells. Upon reaching OD_600nm_ ∼0.8, 0.5µL of culture is spotted on a 150µm thick gel composed of 1% agarose dissolved in the corresponding growth medium used for cultivating cells. Samples were mounted with a coverslip and sealed with parafin. Images were acquired using a Nikon Ti-E inverted microscope (Nikon, Japan) equipped with the Perfect Focus System (Nikon, Japan), a phase contrast objective (CFI Plan Apo Lambda 100X oil NA 1.45), a SemrocK DAPI filter (Ex: 470/20nm Em: 520/20nm), an Orca R2 camera (Hamamatsu Photonics, Japan) and managed with NisElements software (Nikon, Japan). Typically, 9 to 16 fields separated by 200nm were photographed. Samples were illuminated with white light (SpectraX, Lumencor LED) for 200ms at 95% power (target average signal intensity of 20 000 a.u.). GFP was excited with the 470nm led of the SpectraX light source set at 90% power for 800ms.

#### Image analysis

Cell masks were generated from phase contrast images using Omnipose [28]. MicrobeJ o.17 [29] was used to generate fluorescence profiles along the cell body from phase contrast and fluorescence images as well as Omnipose cell masks. Fluorescence profiles (50 bins along the cell length) were normalized by setting the maximal intensity along the cell body to 1, and were then sorted by cell length to generate a demograph (matrix of 50 columns and as many rows). The fluorescence profiles were subjected to principal component analysis to identify the principal axes that could be used to separate groups of cells according to the fluorescent intensities peaking or not at 1/2, or 1/4-3/4 positions along the cell body. The proportions of cells in each category were used to estimate cell cycle period durations using equations 1.

### Statistical analyses

All statistics used throughout the study were performed using Python 3.12. Wilcoxon signed-rank tests was used to analyse potential differences between stes of paired samples. Kruskal Wallis H-test or one-way ANOVA was used to test for effect of a treatment or condition on sample median or mean, respectively, depending upon the validation of the Normality and heteroschedasticity assumptions associated with ANOVA. All statistical analyses are available as a supplementary file (link). .

### Model and parameter estimation

The model described in [7] was translated into a set of python methods bundled into a *Model* class in *bacycle*. These methods were used to generate theoretical cytometry profiles based on 4 parameters (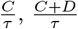, and two parameters required to model a Log-Normal noise, *cv*, and a conversion factor *f*_*c*_ translating fluorescence intensity into genome equivalents. Theoretical or experimental profiles were fitted using the Nested Sampling approach [21, 22] of the *dynesty* python package [23, 30] to sample the joint posterior distribution of parameters. The python package dedicated to the estimation of the cell cycle parameters is based on the dynamic Nested Sampling algorithm [31, 32] included in the *dynesty* package.

To understand the Bayesian approach to fitting observational data, let’s assume an experimental set-up where, given some data *d* and assuming a model *M* parametrized by parameters *θ*, we wish to explain the data using the model. The (posterior) probability that the given data is explained by a set of parameters *θ* of the model *M* can be written:

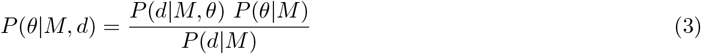

where *P* (*d*|*M, θ*) ≡ ℒ (*θ*) is the likelihood of observing the data given some parameters for a fixed model M. *P* (*θ*|*M*) ≡ *π*(*θ*) is the prior probability of parameters *θ*, based on prior knowledge on the possible ditribution of model parameters, before taking any data.

*P* (*d*|*M*) ≡ *Ƶ*(*θ*) = ∫_Ω(*θ*)_ ℒ (*θ*)*π*(*θ*)*dθ* is a measure of how well the model *M* explains the observed data *d*, taking into account all possible values of the parameters *θ* and their probability distributions. The posterior distribution *P* (*θ*|*M, d*) is our updated knowledge of probable parameter values, informed by the observed data. This is the quantity we are interested in here.

### *bacycle* usage

The *bacycle* Python package requires a Numpy array of single cell fluorescence intensity proportional to DNA amounts. *bacycle* includes utility functions based on *flowio* package 1.4.0 to parse and read *\**.*fcs* files. *bacycle* uses *dynesty* as an established Python implementation of a Nested-Sampling algorithm [23, 30]. When inferring parameters from single cell fluorescence data, this is the *dynesty* output that indicate the evolution of the inference. This dynamic output provides the time elapsed during Nested Sampling optimization.

The following number indicates the number of iterations of the estimation of the model per second. On the same hardware setup we typically reach 200-300 it/sec, and drops below tenth of iterations per second indicate that the inference procedure is hampered, most likely because new meaningful Live points are located in low probability regions of the priors. To circumvent such time delays, it may be useful to use uniform priors to start with instead of using Gaussian or Log-normal priors without prior knowledge on the location of parameter values.

Some inference parameters are directly controllable as input parameters of the *Inference* class, while others are fixed in the code but can be modified if needed in the definition of the class. The parallelization is controlled with the *multiprocessing* package and the number of CPU’s used by the likelihood maximization algorithms can be specified with *ncpus* parameter. The *nchains* parameter specifies the number of parallel chains run by the *emcee* MCMC inference algorithm. Concerning the *dynesty* sampler, the *dlogz init* controls the stopping criterion based on the tolerance on the increment of the evidence Ƶ between iterations, while the *nsamples* fixes the maximum number of iterations. Among fixed parameters (could be changed by modifying the code), the number of Live points for the Nested Sampling algorithm (*nlive* parameter of *dynesty*) is set to 128, and could be increased to help inferences that converge too slowly. Similarly, the type of move is set for *emcee* as a combination of differential evolution moves (80% DEMove() [33], 20% DESnookerMove [34], which uses a Snooker updater for the differential evolution adaptative algorithm).

### Code and data availability

*bacycle* is available as a Python package. Data are publicly available on the Zenodo archive under the unique identifier DOI:10.5281/zenodo.18556582.

## Results

### The standard rifampicin run-off approach fails for many *E. coli* isolates

Flow cytometry has had a major impact on cell-cycle studies of mammalian cells. However this method has not been widely used in microbiology, probably because of the bacterial cell cycle specificity and the lack of automated and dedicated analysis tool. Nevertheless, a robust procedure to acquire DNA amount per bacterial cell by flow cytometry has been established over the last 2 decades the last century [18, 35]. Following this procedure, we measured DNA amounts by FRET with a combination of Mithramycin A and ehtidium bromide. Mithramycin A binds the small groove of DNA at G or C dimers [36]. We emphasize the importance of the Mg^2+^ ions added to the staining buffer to lower the solubility of DNA fragments and prevent loss of DNA from cells [37].

Under fast growth conditions, distributions of cellular DNA content do not appear to be highly structured (Figure 1, gray curves). The distribution of DNA amount per cell quickly separates in 2 distinct peaks after rifampicin and cephalexin treatment corresponding to cells with n and 2n genome equivalents, as observed in the laboratory *E. coli* K12 MG1655 strain (Figure 1A). However, we find that the antibiotic treatment does not result in a crisp separation of 2 sub-populations based on DNA content for other model *E. coli* strains such as the UPEC *E. coli* CFT073 and *E. coli* strain E22 (Figure 1B, C). These strains are neither resistant to rifampicin nor cephalexin and cell growth is readily arrested by the addition of rifampicin alone (Figure S1). In the case of *E. coli* CFT073, the antibiotic treatment initially inhibits the initiation of DNA replication while ongoing replication rounds are terminating, as indicated by the formation of the 2 peaks at the earliest time point after treatment (Figure 1B, compare before and after 30min of antibiotic treatment). However, the proportion of cells with intermediate DNA content (between n and 2n genomes) remains relatively stable and never transition toward 2n genome equivalents. Thus, it appears that ongoing replication is also inhibited quickly after treatment and do not terminate, even when the treatment lasts for more than 7 generation times (Figure 1B). Strikingly, the antibiotic treatment has virtually no effect on the distribution of genomic content per cell of *E. coli* E22 cells (Figure 1C), despite the growth arrest induced by the addition of rifampicin to a culture of E22 cells (Figure S1). Thus, the pharmacological treatment adapted for replication run-off in the laboratory *E. coli* strains appears to inhibit DNA replication altogether in *E. coli* E22 cells, which results in a snapshot of the distribution of cellular DNA content similar to the distribution before treatment (Figure 1C). This differential effect of the antibiotic treatment on DNA replication prevents replication run-off and thus, the estimation of C and D periods from such cytometry profiles.

**Fig 1.**
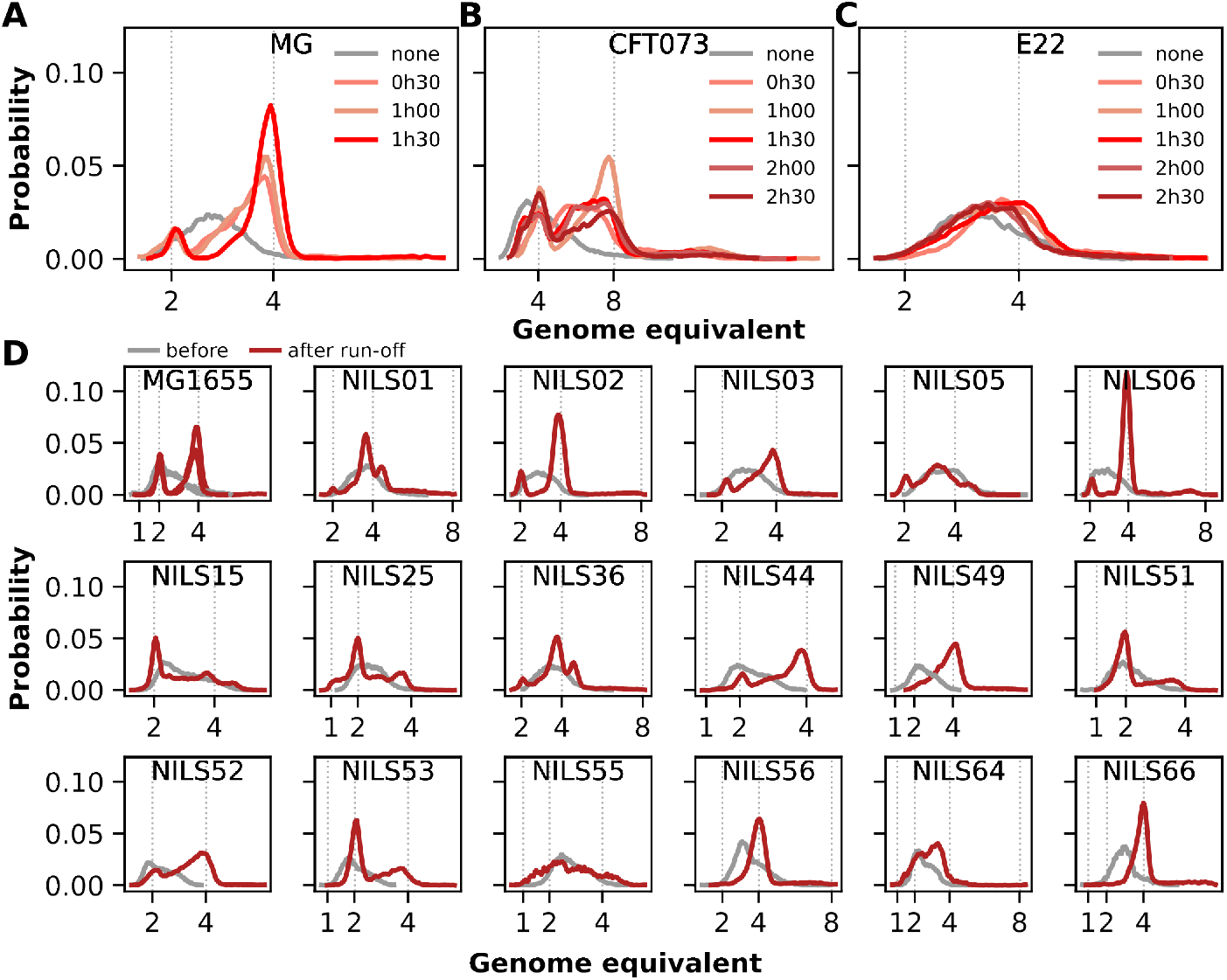
Effect of rifampicin-cephalexin treatment on DNA replication for multiple *E. coli* strains. A-C) Distributions of DNA amount per cell over a time course experiment showing replication run-off over time, from 30min (light red) to 2h30 (dark red curves) for A) K12 MG1655, B) CFT073 and C) E22. Gray curves illustrate the probability density distribution of cellular DNA content before treatment. D) Distributions of DNA amount per cell for end-point replication run-off experiments for 17 NILS strains distributed accross the *E. coli* species. Rifampicin was added at a final concentration of 150*µ*g/mL and cephalexin at 10*µ*g/mL. Red curves represent cellular DNA content distributions after antibiotic treatment.

To test how common is this inhibitory effect of rifampicin on ongoing replication forks, we attempted to perform replication run-off experiments on a set of *E. coli* strains representative of the *E. coli* species. To this end, we selected a set of 17 strains from the Natural Isolates with Low Subcultures (NILS) collection [38] according to their genomic divergences so as to cover as much as possible the genetic diversity of the *E. coli* species. While the rifampicin-based replication run-off experiment consistently leads to separated peaks for the laboratory strain *E. coli* K12 MG1655, only 4 out of 17 NILS strains presented typical run-off profiles, with well separated peaks at *n* and 2*n* genome equivalents Figure 1D, NILS02, NILS06, NILS56 and NILS66). In the cases of NILS56 and NILS66, the distribution of DNA amount per cell after replication run-off tends to form a single peak around 4 genome equivalents, albeit a bit asymetric toward lower DNA content and a wider than the 4 genome equivalents observed for NILS02 or NILS06. For these 2 strains, the replication run-off seems to be as efficient as for the reference strain, but with a duration of the C+D period very close to twice the generation time. In these conditions the initiation of DNA replication coincides with cell division. As a result, nearly 100% of cells present 4 genome equivalents. For the remaining 13 NILS strains, the flow cytometry profiles are reminiscent of the run-off profile of *E. coli* CFT073 cells. It appears that in most cases, ongoing DNA replication cycles are more or less quickly arrested and a good proportion of cells with non-integer genome equivalents remains in the population, even after 6 to 7 generation times of antibiotic treatment.

### bacycle: a Python package for an objective estimation of cell cycle parameters

Building up on the pioneering work from Skarstad and colleagues [18, 20] and from Hansen and colleagues [19], we developed a Python package dedicated to the objective search for the best parameters reproducing cytometry profiles of cellular DNA content per cell without the need for drug treatment. *bacycle* is built on the cell cycle model proposed in 1968 [7] and proposes 2 established Monte Carlo approaches to fit model parameters. The amount of fluorescence signal detected for each cell, after gating out other particles, is the only input needed - provided as a NumPy array [39].

#### Model and methods

The bacterial cell cycle model proposed by Cooper and Helmstetter [7] is used to calculate the probability distribution of DNA amount per cell in an asynchronous population according the duration of C and D periods, that are considered fixed for all cells of the population as cells are hypothesized to be in a dynamical steady state during exponential growth. These 2 biologically relevant parameters (cell cycle period durations) need to be combined with 2 technical parameters to generate a theoretical flow cytometry profile for comparison with experimental data: a conversion factor, *f*_*c*_, translating DNA amount into fluorescence units and a noise parameter, *cv*, that models experimental uncertainty in evaluating the fluorescence signal from any given cell.

*bacycle* includes 2 different optimization procedures, that can be specified with the *sampling algorithm* input parameter of the *bacycle* inferencer, to estimate parameter values in the relatively low-dimensional parameter space (4d).

i. Dynamic Nested Sampling with *dynesty* [23, 30] *(sampling algorithm=‘dynesty’*). This is the main approach outlined here.
ii. Markov Chain Monte Carlo (MCMC) approach with *emcee* [40] (*sampling algorithm=‘emcee’*). This affine invariant MCMC implementation requires an initial guess for parameter values (input parameter *theta0*).

#### Model parametrization - setting the priors

Both algorithms optimize a set of 4 parameters to match the input data: 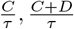, *cv, f*_*c*_ (Figure 2A). The first 2 are the biologically relevant parameters that we want to fit and can only be constrained to the range of biologically relevant values. By default, the relative duration of the C period is constrained to the 0.4, 2.6 range, meaning that the duration of the C period initially varies from 40% to 260% times the generation time (*τ*) (from poor to rich growth conditions). The relative duration of the C+D period is searched by default in the [0.4, 4] range, that is from a situation where the sum of the C and D periods represent 40% to 400% of the generation time. These constrains may be changed by explicitly defining the form and range of probable values of the priors on these 2 parameters (*i*.*e*., setting *model*.*distributions[‘CD tau’]* as a uniform distribution between 0.4 and 4, or any pair of values – see *bacycle* [help page](https://src.koda.cnrs.fr/genome-dynamics/bacycle)).

**Fig 2.**
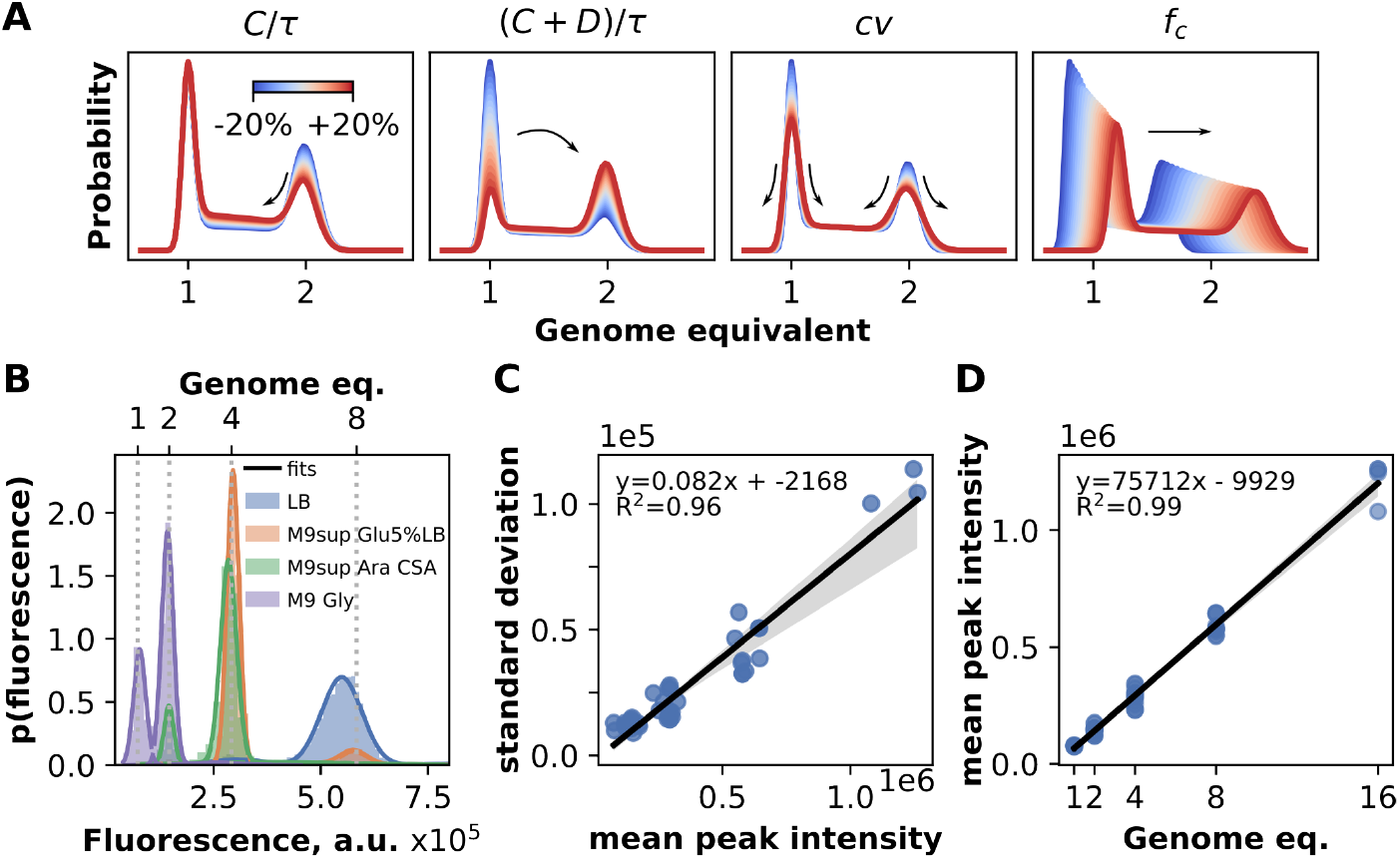
Noise calibration using rifampicin run-off experiments. A) Example of simulated cytometry profile alterations induced by changing a single parameter value by 20% around its original value for an initial parameter set 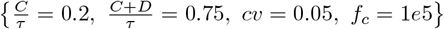. Increasing 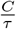 values without changing 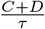 reduces the duration of the D period and increases the fraction of cells actively replicating to the detriment of the fraction of cells having completed this replication round. As 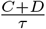 values increase, the earlier replication initiates relative to cell birth and the average amount of DNA per cell increases. Increasing *cv* values further smooth the signal. Increasing *f*_*c*_ values shifts the profile to the right (more DNA per cell). B) Representative distribution of cellular DNA content from an asynchronous population of cells treated with cephalexin and rifampicin colored by growth conditions are fitted with a Gaussian mixture model so that each peak is captured by one Gaussian component. The Gaussian mixture components add up to the global fit to the data (colored solid lines). The corresponding genome equivalents are illustrated on the second x-axis represented above the graph. C) Scatter plot of the mean and standard deviation of the 2 dominant peaks for each replication run-off experiment carried out on multiple *E. coli* strains, under multiple growth conditions. Each blue circle represents one peak. The linear regression (black line) and its associated 95% confidence interval (gray shaded area) illustrate the linear dependence of the standard deviation of each peak on the mean fluorescence of each peak (e.g., constant *cv*). The slope of this best linear fit is used as an estimation of the noise parameter *cv* (0.082 +/-0.007 - 95% confidence interval). D) Scatter plot of the mean fluorescence per chromosome from replication run-off experiments on the strain *E. coli* K12 MG1655 grown in different growth media. The linear regression to the data and its 95% confidence interval is illustrated with the black line and shaded grey area, respectively. The thick black line represents the regressionline to data and the gray shaded area represents the fit 95% confidence interval. The slope of this best linear fit is used as an estimation of the *f*_*c*_ parameter (75712 +/-3034).

All other parameters being constant, each parameter influence the shape and scale of the distribution of cellular DNA amounts in its own way (Figure 2A). Shifting 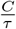 without changing 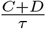 modifies the fraction of cells undergoing replication at the expense of the fraction of cells in the D phase. As a result increasing 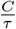 up reduces the fraction of cells with the highest DNA amounts and inflates the fraction of cells with lower DNA amounts. Shifting 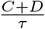 up or down (without changing 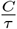) modifies the fraction cells in the D phase and expands or shrinks, respectively, the right side of the distribution. Increasing the *cv* erodes the distribution, while the conversion factor *f*_*c*_ shifts the location of the distribution (shift along the x-axis).

Earlier work emphasized the importance of a proper noise model to capture the dependence of the measurement error with the fluorescence level of a particle. Both seminal studies on the topic noted that the detection noise was characterized by a constant coefficient of variation (standard deviation divided by the mean, 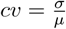) [18, 19]. Hence, the standard deviation for the detection of a particle type is proportional to its mean fluorescence, modeled as Log-Normal noise. To characterize the type of noise associated with our experimental conditions, we opted for the quantification of the variations of fluorescence detections associated with bacterial cells containing integer number of chromosomes for two reasons. This approach allows for the noise characterization (i) using the same experimental procedure as for experiments and (ii) within the relevant range of fluorescence units detected in experiments. Using the laboratory strain *E. coli* K12 MG1655, amenable to run-off experiments, we compiled cytometry profiles under various growth conditions encompassing situations with cells containing from 1 to 8 chromosomes per cell after replication run-off (Figure 2B). As expected, the cytometry profiles are dominated by 2 major peaks corresponding to the number of origins at birth and at division (twice as much DNA at division). A third peak may appear in replication run-off experiment because antibiotic treatments do not yield perfect inhibition of DNA initiation or cell division. Considering the population of cells with *n* or 2*n* chromosomes as homogeneous, we modeled each peak as a Gaussian distribution and estimated its mean and standard deviation using a Gaussian mixture model. Figure 2C illustrates the linear dependence of the standard deviation of peaks with the mean fluorescence of each peak, or equivalently, that in our experimental set up we also find the noise to be characterized by a constant coefficient of variation. We estimated the *cv* to be close to 8% (0.082 +/-0.007 - 95% confidence interval) in this set of flow cytometry experiments. We note that this value is larger than the one estimated from calibration experiments carried out with fluorescent beads, which is close to 3% (Figure S2) and corresponds to the precision reported by the cytometer constructor (CytoFlex, Beckman Coulter). The higher noise detected with our experimental set-up suggests that the experimental noise is not dominated by instrumental noise (*i*.*e*., fluorescence detector noise).

The same set of replication run-off experiments can be used to experimentally derive an estimation of the last parameter *f*_*c*_, that translates fluorescence units into genome equivalents. Replication run-off experiments generates populations of cells with integer number of chromosomes as illustrated by fluorescence peaks and vertical dashed gray lines in Figure 2B. The mean fluorescence associated with integer chromosome values is well described by a linear model, indicating that the fluorescence signal is proportional to the DNA content (Figure 2D).

In the case of the *E. coli* K12 strain MG1655 and under the specific labeling protocol used in this study, this proportionality factor is estimated to be close to 74 000 fluorescence units for 1 chromosome equivalent (73 897 +/-3 034 - 95% confidence interval) (Figure 2D). Note that this relationship depends on the concentration of dyes and on the specific dye batch used, therefore, a control sample should be run at each cytometry acquisition for calibration. Other methods could be used to obtain the same information. While replication run-off experiments may be carried out in some cases with chloramphenicol or serine hydroxamate, these treatments are not stringent enough to properly estimate the C+D period. However, these treatments may eventually generate population of cells with integer number of chromosomes and be useful to estimate the fluorescence amounts per chromosome. Alternatively, and most importantly irrespective of strains sensitivity to drug treatments, bacterial strains can be grown under slow growth conditions. When the C+D period is shorter than the generation time *τ*, the population of cells in either B or D periods contribute to sharp peaks corresponding to 1 and 2 chromosomes per cell. These peaks provide a general mean to estimate the conversion factor *f*_*c*_ for a given strain under specific experimental condition (*e*.*g*., labeling procedure, fluorescent dyes concentrations).

The model considered is therefore built on 2 biologically relevant parameters and 2 technical parameters that can be constrained based on experimental data. Based on experiments presented above, the level of noise (*cv*) can be adjusted around a value of 0.08, while the conversion factor can be constrained around the experimentally derived value of ∼74 000 fluorescent units (for this set of acquisitions for *E. coli* K12 MG1655 and in this experimental setup).

### Finding unique solutions with experimentally derived parameter constrains

When multiple sets of parameters may reproduce the same cytometry profile equally well, the model is deemed to be non-identifiable, in the general sense. However, these parameter sets may be far apart in the parameter space, and may therefore correspond in reality to very different situations. In this case, some knowledge about the experimental conditions may be sufficient to exclude all but one of the estimated parameter sets as a reasonable solution.

The Cooper-Helmstetter model is not generally identifiable, leading to the fact that multiple parameter sets can describe the same cytometry profile. Figure 3 illustrates the multiplicity of solutions for a pair of biological parameters 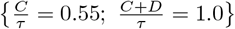. Alternative mathematically satisfying solutions appear for biological parameter values raised or reduced by one unit. In the case illustrated in Figure 3A, echoing solutions (purple areas) occur at 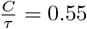 and 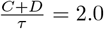 or 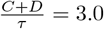. The simulated cytometry profiles associated with these parameters correspond to very different DNA amounts per cell (Figure 3B), but share the same shape and are indistinguishable when multiplied by the proper *f*_*c*_ parameter (Figure 3C).

**Fig 3.**
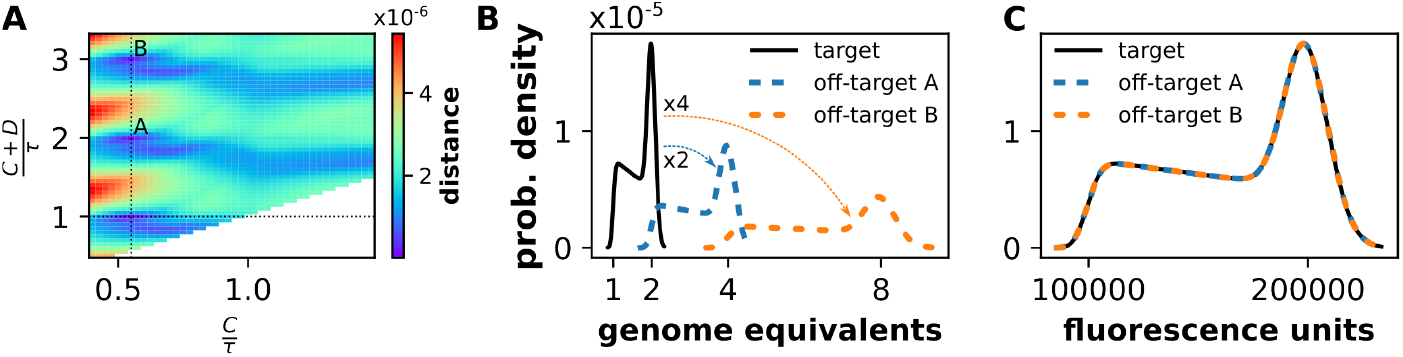
The Cooper-Helmstetter model is not generally identifiable. A) Matrix of distances between the target probability distribution function (pdf) simulated with parameters [0.55, 1.0, 0.05, 1.10^5^] with the pdf generated with other values for the first two parameters (biological parameters in *x* and *y* -axes). The distance (color coded) is calculated as the Wasserstein distance and is independent of technical parameter *f*_*c*_. Three zones of short distances appear, indicated by the crossing of the 2 black dashed lines (true parameter values) and by the characters A and B (2 off-target solutions). Parameter space where 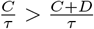 is not accessible by definition (white area). B) Illustration of the 3 probability density distributions of genomic content per cell corresponding to the 3 parameter sets. The target distribution (thick black line) is shifted to higher DNA amounts by increasing 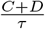 from 1 to 2 (off-target A – dashed blue line) or to 3 (off-target B – dashed orange line). C) Once shifted and normalized by the proper conversion factors (*f*_*c*_ values divided by 2 and 4, respectively), the 3 distributions are indistinguishable.

Echoing solutions typically appear by shifting the biological parameter values by an integer value (see Figure S3 for a different parameter set and higher number of alternative solutions). Echoing solutions seldom are as good fits as the true parameter set, and in most situations, even without constraining down the *f*_*c*_ parameter, the Nested Sampling algorithm identifies the true parameter set as the most probable solution.

Constrains on parameters are necessary to make the model locally identifiable. Specifically, the *f*_*c*_ parameter (conversion from fluorescence signal to DNA amount) needs to be estimated within a 20% error range to avoid identifying multiple solutions fitting a given experimental cytometry profile. Constraining the biological parameters within their range of biologically relevant values accelerate the convergence of the optimization algorithm toward a solution. With the prior constraints described above, *bacycle* typically finds a solution in 1–2 minutes on a single CPU. While *dynesty* supports parallelization, the reduction in completion time does not scale linearly with additional CPUs. Nevertheless, using more CPUs can still accelerate the computation.

### Bias and precision from simulated cytometry profiles

Experimental and biological noises alter ideal measurements and estimation procedures must be robust to noise levels. But it is also important to verify first that the optimization procedure does not introduce any bias, or is not highly imprecise by design. *bacycle* is based by default on a Bayesian inference algorithm (Nested Sampling). As such, parameters are estimated from the posterior distribution, that is the probability of observing parameter values based on the data. The marginalized posterior distribution for each parameter is therefore precisely a measure of our degree of certainty about each parameter value. Typically, an inter-quantile range of these marginalized posterior distribution is used to define a *credible interval*. Note that credible intervals and confidence intervals are 2 different notions. While the Bayesian credible interval actually provides an estimation of the marginal probability that the distribution should take a given value, the frequentist confidence interval proposes boundaries within which the fixed observed distribution value would fall if we were to repeat the sampling many times. Thus, a credible interval covering a fraction *f* of the posterior distribution is by construction an interval in which the parameter value is expected to be included with probability *f* . We chose to use median credible intervals of probability *γ*, with the 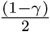 and 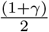 quantiles of the posterior distribution as boundaries (*i*.*e*., with *γ* = 95% the credible interval is bounded by the 2.5% and 97.5% quantiles of the posterior distribution – see Figure 4A).

**Fig 4.**
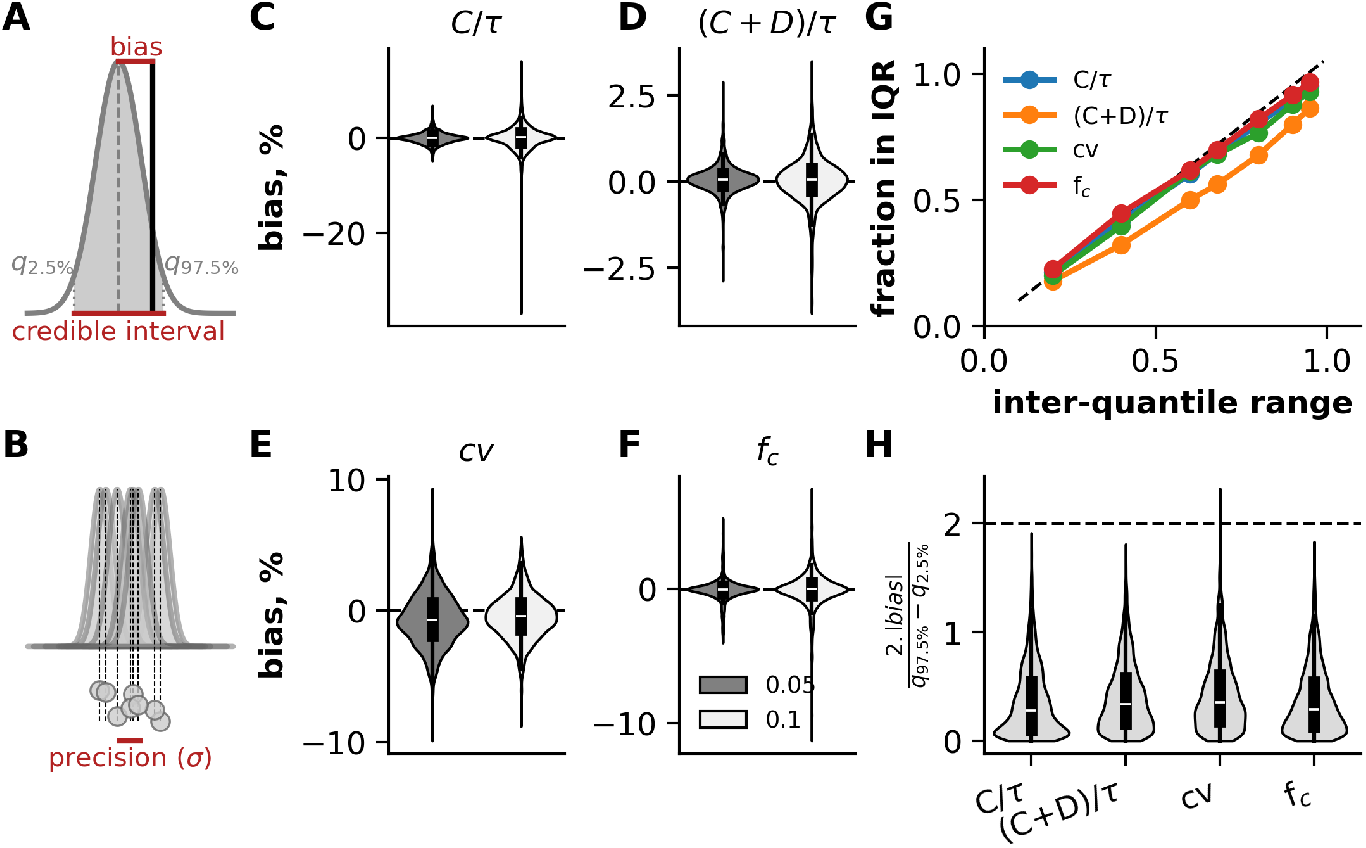
Low bias of Nested Sampling inferences from inferences from simulated, theoretical cytometry profiles. A) Illustration of bias and credible interval from a likelihood distribution. The gray curve shows a hypothetical likelihood distribution sampled through the Nested Sampling procedure. The even-tailed 95% credible interval encompasses 95% of the posterior probability distribution from the 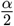 and 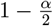 quantiles, with *α* = 0.05. The relative bias was calculated as the deviation of the median of the posterior probability distribution from the target parameter value symbolized as the vertical black line: 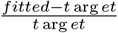. B) The precision of the method was calculated as the standard deviation of multiple (n=50) inferences on theoretical cytometry profiles generated from the each parameter each parameter set. C-F) Violin plots in each panel show the distribution of the relative bias calculated from all repeated inferences for the 4 parameters. Cytometry data were simulated over a range of parameter sets for both biological parameters and 2 noise levels (cv=0.05 or 0.1). Positive (negative) bias values indicate an overestimation (underestimation). See Figure S4 for a representation of the bias for each pair of biological parameter values. G) QQ-plots for the 4 parameters illustrate the correspondence between the credible interval range (x-axis) and the fraction of the credible intervals containing the target value (y-axis). The x=y diagonal is indicated as a black dashed line. H) The violin plot illustrates for each parameter the distribution of the ratio between bias and the half-range of the 95% even-tailed credible interval. 95% CI is expected to extend to 2 standard deviations from the mean and this threshold is indicated with an horizontal dashed black line. 95% of the absolute bias values are expected to fall below this threshold.

We verified that Nested Sampling algorithm used on data simulated with the Cooper-Helmstetter model properly estimate parameters used to simulate the cytometry profile in the first place, without bias (Figure 4A) and with the highest possible precision (Figure 4). By repeating the data simulation and optimization procedure 50 times for each set of a range of parameter values, we showed that the Nested Sampling algorithm leads to virtually unbiased estimations as shown by the violin plots centered on 0 in Figure 4C-F), except for a slight under-estimation by 1 to 2% for *cv*, the noise parameter. While the 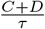 seem to be robustly and precisely estimated in an unbiased manner regardless whether we included 5 or 10% of noise in the simulated data (bias values never extend beyond 3%), other parameters, and in particular 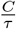 can, in some rare instances be more strongly biased (beyond 30% under-estimation, which would be biologically significant). It is therefore important to note that *bacycle* provides precise, unbiased estimations of 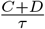, even if it can be less precise for the estimation of 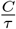 (Figure 4C). This differential effect on the precision on the 2 biological parameters is enhanced as the true 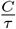 and *cv* value increase (Figure S4).

We further verified that the *γ*-credible intervals included true parameter values at the expected frequencies – that is that the *x* % credible intervals included true parameter values with the expected frequency *x*, as indicated by the qq-plot in Figure 4G. The fraction of estimations falling within the inter-quantile at a given level follows the expectation indicated by the *y* = *x* diagonal. The slight deviation for the 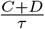 parameter (orange curve) indicates that the width of the posterior distribution is systematically narrower than expected, with the orange line positioned marginally below the diagonal. On average, this results in an 8.9% underestimation, with inter-quantile ranges failing to include the true value approximately 9% more frequently than anticipated. Therefore, the credible intervals can be interpreted as uncertainty levels about the estimated parameter values, but a slight bias for the 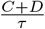 parameter. Furthermore, we verified that for each of the 4 parameters and regardless of the parameter values, the scattering of biases over 50 repeated inferences is smaller than the half size of the 95% credible interval (Figure 4H). Low ratio values suggest that the uncertainty level measured with the Nested Sampling algorithm is typically larger than the variation of the bias between estimations. In other words, the bias remains small relative to the interval width, which serves as an indicator of good inferential quality.

Overall, the inferences produced by *bacycle* are precise and unbiased by design, especially for the 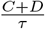 parameter. In addition, we could verify that the credibility intervals produced by *bacycle*, via the Nested Sampling algorithm encoded in *dynesty*, can be interpreted as uncertainty levels on parameter estimations. These results indicate that the inference procedure properly explores the parameter space according to the Cooper-Helmstetter model and do not introduce strong bias or high variability by design.

### Estimations of 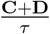 with *bacycle* on *par* with drug-mediated replication run-off bacycle estimates compares with published data and replication run-off experiments

Turning to experimental data, we asked how our parameter estimation procedure fares when compared with the state of the art pharmacological run-off experiment. We collected *E. coli* K12 MG1655 cells grown in 5 different growth conditions before and after the addition of rifampicin and cephalexin for a period of time spanning at least 3 generation times. We then estimated for each growth condition the 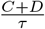 parameter from the run-off sample using the fraction of cells in the *n* and 2*n* chromosome peaks, or *bacycle* and the nested sampling algorithm for the untreated cells. We also compared our results to previously published data for the same *E. coli* K12 MG1655 strain. Figure 5A shows that the 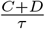 estimates from the untreated cells are very close to the values derived from the state of the art replication run-off experiments (red versus orange dots) and are comparable to the values published over 2 decades for this same strain by multiple research groups (gray versus colored symbols). In our hands, both experimental approaches lead to similar levels of reproducibility, with coefficients of variation (*µ/σ*) below 10% for all 5 growth conditions (Figure S5A), covering a range of generation times from 24 to 140 min.

**Fig 5.**
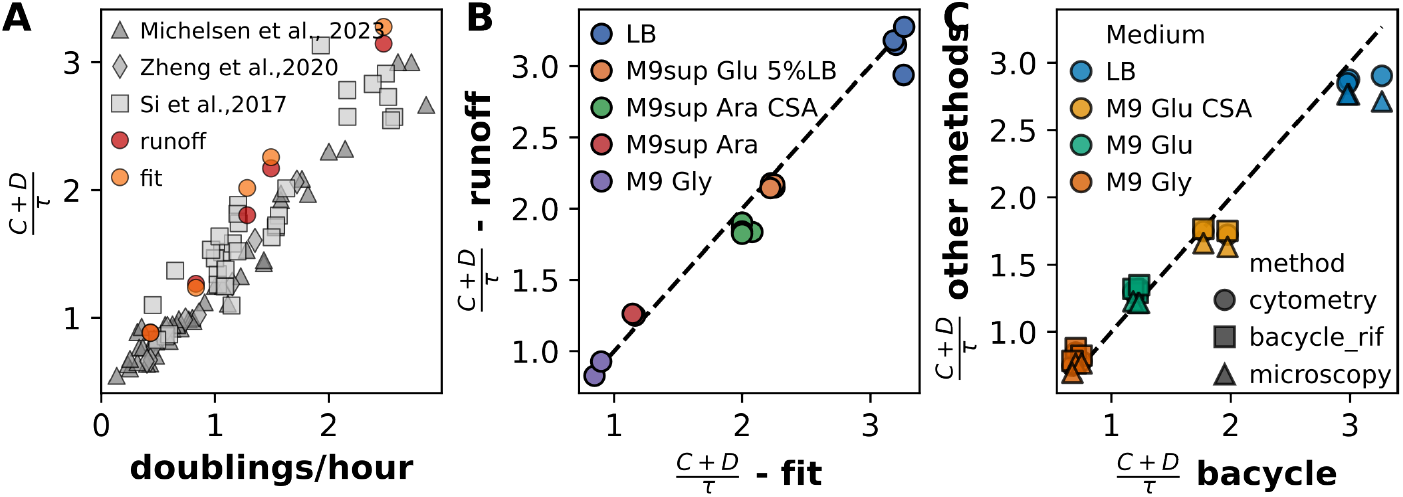
Fitted values compares with published data and state-of-the-art replication run-off method. A) Comparison of 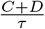 values from the fitting procedure, replication run-off experiments on *E. coli* K12 MG1655 and with previously published data. Scatter plot showing the dependence of 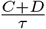 on growth rate. Red and orange circles represent replication run off derived values and fitted values generated in this study, respectively. Gray symbols represent previously published data. B) Comparison of 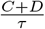 values fitted with *bacycle* (x-axis) and estimated by replication run-off (y-axis) on the same cultures of *E. coli* K12 MG1655 grown in different growth media (color coded). C) Scatter plot comparing 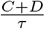 estimated with different methods (different markers) obtained from the same cultures of *E. coli* K12 MG1655 *dnaN::gfp-dnaN* grown in different growth media (color coded). The black dotted *y=x* diagonal line on panels B) and C) indicates the expectation for a perfect agreement between methods.

We further verified the agreement between *bacycle* and replication run-off derived estimates by comparing 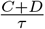 results obtained from cells sampled from the same culture. In these experiments carried out in 5 different growth media, cells are sampled in exponential phase just before and after the rifampicin and cephalexin treatment. These paired samples are used to estimate 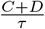. Figure 5B shows that both methods lead to estimates that are very close to one another (Pearson r= 0.98). In fact, the 2 methods are statistically biased with respect to one another (Wilcoxon signed-rank test on 18 pairs, p-value=0.012, effect size 0.43 – see Supplementary Information). But the slight positive bias of *bacycle* over run-off (without implying any priority between the 2 methods) is too small to be biologically and practically relevant. The median of the differences between pairs of estimations is 0.075 in 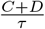 unit, which is in the range of the experimental variability mentioned above and measured over a larger set of unpaired *bacycle* estimations. Standard deviations range from 0.052 for LB, to 0.084 for M9sup Ara without rifampicin, and from 0.022 for M9sup Glu 5%LB to 0.146 for LB (Figure S5A).

The conversion factor *f*_*c*_ from fluorescence to genome units needs to be constrained to favor identifiability, but the noise *cv* parameter was left largely unconstrained, with a uniform prior ranging from 2 to 25%. While replication run-off samples lead to similar *cv* estimates as indicated earlier (Figure 2C), around 6-10%, untreated samples were fitted with higher *cv* values (15-25%, Figure S5B). The slow growth condition (M9 Glycerol) is the only condition where the *cv* is estimated at the same level as for replication run-off samples. The over-estimation of the noise suggests that in cell cultures growing at fast growth rates, there is another source of variability that is not modeled in the Cooper-Helmstetter model. This source of noise is likely of biological origin because it disappears when the same cell cultures are treated with rifampicin and cephalexin.

We sought to further compare *bacycle* and replication run-off methods by fitting cytometry profiles after replication run-off (the same ones as for the standard method) and with a third approach. To do so, we evaluated cell cycle period durations using fluorescence microscopy on a strain expressing a fluorescent fusion with the *β*-clamp loader protein DnaN from its native locus (*dnaN::gfp-dnaN* [25]). The relative timing of cell cycle events can be inferred from the fraction of cells that have passed a specific cell cycle stage [41, 42]. This relative timing inference method has been used and tested by multiple groups [43–45]. Here, we sought to define the relative timing of C+D (*i*.*e*., 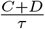) by identifying the fraction of cells with GFP-DnaN fluorescent foci, indicative of active replisomes. We performed the same flow cytometry analyses as before (with and without rifampicin + cephalexin treatment) on the same cell cultures that were imaged under the microscope (Figure S5C-F and G-J). The presence of fluorescence spots at mid-nucleoid (*e*.*g*., mid-cell for non-overlapping cell cycle, or 1/4 - 3/4 positions for 2 concurrent cell cycles) indicate ongoing replication. The analysis of the fluorescence intensity along the cell body over an asynchronous population of cells can therefore reveal the proportions of cells undergoing replication (Figure S5G-J). The comparison of *bacycle* 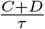 estimations with replication run-off and fluorescence microscopy shows that all 3 methods are in good agreement (Figure 5C), with small systematic biases between the methods. Notably, the two most closely related methods both utilize cytometry profiles following replication run-off: the current state-of-the-art approach (gating cytometry profiles post-run-off) and the application of *bacycle* to the same cytometry data (post-run-off). These 2 methods deviate from one another by 0.026 unit (mean, n=10). Running *bacycle* on cytometry profiles from untreated cells produces results that deviate slightly from the above-mentioned methods, cytometry and *bacycle* on run-off profiles, by 0.03 and 0.12 unit, respectively. In this *E. coli* K12 MG1655 derivative, 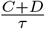 estimations are not significantly different from the standard cytometry method (2-sided Wilcoxon signed-rank test on 10 pairs of values, p-value=0.72, effect size of 0.076 – see supplementary files for a complete statistical report). But this global average mitigates differential biases across the different growth conditions. Differences with *bacycle* estimations tend to be greater as growth conditions improve. The mean deviation with the standard replication run-off method increases up to 0.14 (M9 Glu CSA, n=2) and 0.23 units (LB, n=3). These deviations remain small (*<*10% of the mean) and practically insignificant, but suggest a tendency that is even further noticeable with miroscopy-based estimations that are smaller than *bacycle* estimations by 0.23 units in M9 Glu CSA (n=2) and 0.33 units in LB medium (n=3) (see (Figure 5C)). The introduction of the traductional fusion GFP-DnaN seem to partly impair the synchronicity of the firing of replication cycles, as indicated by the secondary peaks on rifampicin-cephalexin treated samples on flow cytometry profiles (Figure S5C-F), black curves), at 3, 5 or 6 genome equivalents (different from 2^*n*^). These extra peaks make it difficult to properly estimate 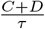 because it is unclear whether each extra peak should be associated the n or 2n genome equivalent fraction. In the same line of argument, the analysis of GFP-DnaN fluorescence profiles along the cell suffers from this very same effect and from the difficulty to clearly associate fluorescence profile to a cell cycle stage (see methods).

#### Estimation of 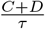 does not depend on the precision on 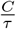

As mentioned earlier, the alterations of the distribution of DNA amount per cell induced by changes in 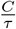 are subtle as growth rate increases (Figure 2A) and this parameter is likely to be most affected by the level of noise. Simulations showed that 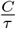 is the most sensitive parameter to noise as indicated by the widest spread of biases. While 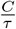 is precisely estimated from simulated profiles, rare improper estimations can deviate by more than 30% from the the target value when noise increases (Figure 4C, see also Figure S4). However, this imprecision on the replication period estimation did not bear any effect on the bias and precision of the 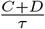 parameter estimation (Figure S4). We sought to verify that this property applies to experimental data. We experimentally measured 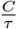 by digital droplet PCR (ddPCR), a technique that does not rely on any drug treatment and that is amenable to any bacterial strain with a sequenced genome. Quantitative PCR have been widely used to measure *ori* to *ter* ratio [46–48], which is related to 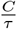 according to the relation:

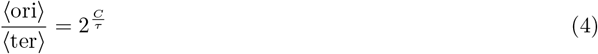

where ⟨ori⟩ and ⟨ter⟩ refer to the average number of origins and termini of replication.

Out of the 5 media tested (see Material and Methods and Figure 6A), we typically obtained a precision under 10% for the estimation of this *ori* to *ter* ratio with ddPCR, with a coefficient of variation close to 3% for cells grown in LB, and up to 7.2% in M9 medium supplemented with glucose and 5% of LB medium.

**Fig 6.**
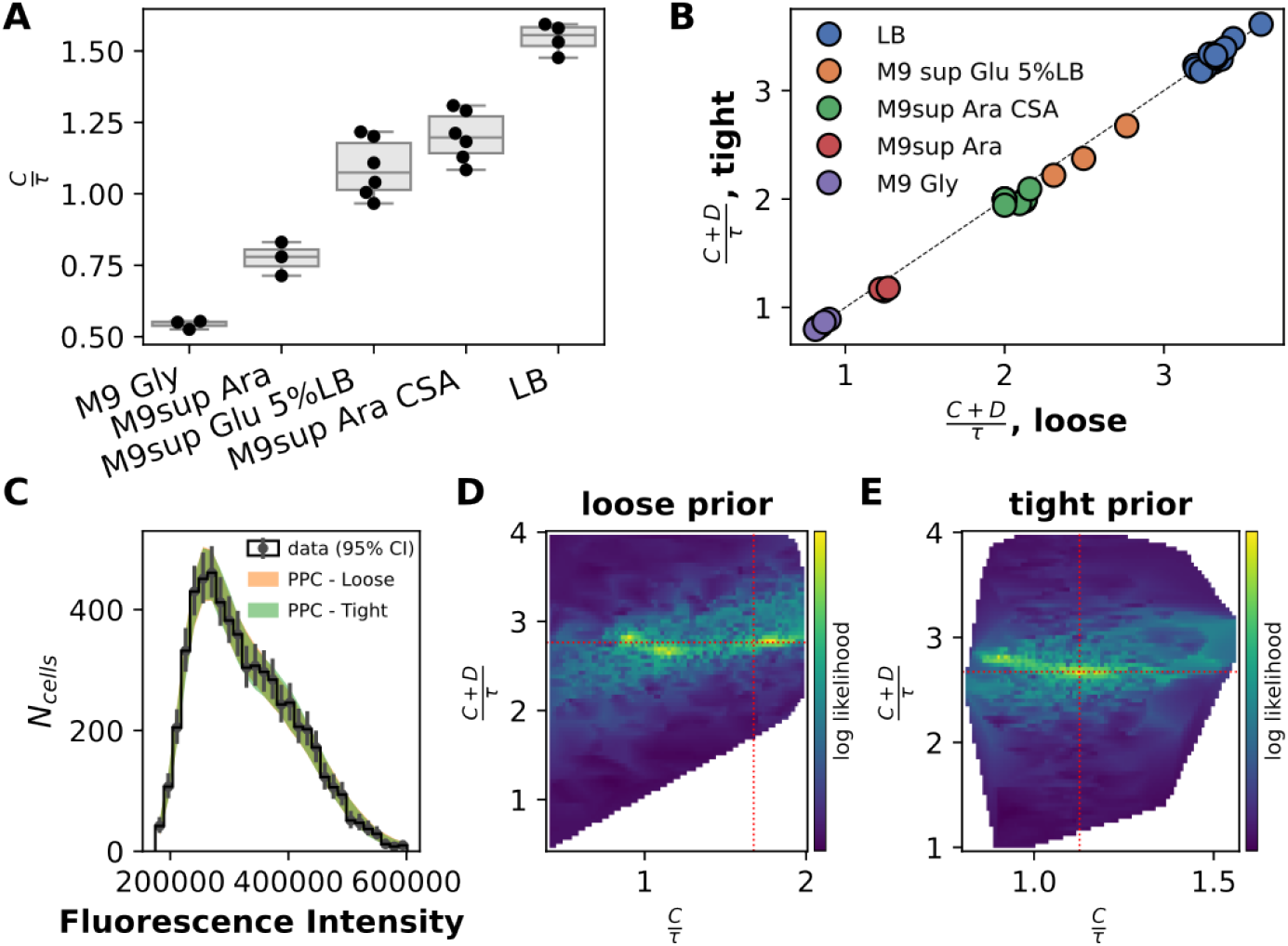
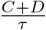 estimation is largely independent from the prior constraint on 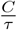. A) Box plot of the 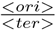 ratio for the *E. coli* K12 MG1655 strain in different growth conditions (growth media). Each black dot represent an individual independent estimation of 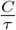 by ddPCR. The boxes and whiskers show the quartiles and extrema of the set of points. B) Scatter plot of the estimation of 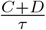 with loose 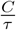 prior – x-axis, uniform over [0.4;2.5] – or tight 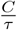 prior – y-axis, Log-Normal with mean and standard deviation based on ddPCR results presented in panel A. Each dot represent the fit with both types of constraints for a given cytometry profile. All cytometry data were generated with *E. coli* K12 MG1655 cell samples, grown in different growth conditions (color-coded by growth media). C) Posterior predictive check (PPC) of both models (with tight or loose constraint on 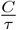) illustrated in panels D) and E). The black line illustrates the distribution of the data, with the error bars representing the 95% confidence interval for each bin. The color shaded areas show the 95% credible interval around the distribution generated from the model with a tight (green) or loose (orange) prior on 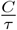). PPCs analyze the degree to which data generated from the model deviate from data generated from the true distribution. D) Log likelihood surface (color coded) explored during the estimation of the model parameters with *bacycle* with a loose prior on 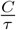 (uniform over the range [0.4, 2.4]) in the 2D space of the cell cycle parameters. The estimated parameters are illustrated with horizontal 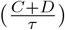 and vertical 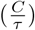 dashed red lines. The Log-likelihood shows 3 maxima (yellow areas) over this parameter space. While 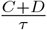 is roughly the same for all 3 minima, the 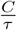 is improperly estimated because of the contribution of all 3 minima and the estimated parameter do not correspond to a true maximum. E) Log likelihood surface (color coded) explored during the estimation of the model parameters with *bacycle* with a tight prior on 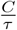 (log-normal with parameters *µ* = *log*(1.1), *σ* = 0.1) in the 2D space of the cell cycle parameters. The estimated parameters are illustrated with horizontal 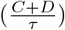 and vertical 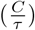 dashed red lines. Both cell cycle parameters are well estimated and correspond to the unique log-likelihood maximum.

We used the experimental measurements of 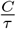 to introduce a tight constraint on the estimation of the 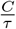 parameter under the form of a Log-Normal prior characterized by the means and standard deviations of 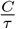 values measured experimentally. We compared the values of 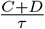 with tight (ddPCR derived Log-Normal prior) or loose (uniform prior) constraint on 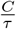. The correlation between these two sets of 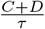 estimates is close to 1 (r*>*0.99, Figure 6B). But statistically, applying a tight constraint introduces a negative bias on 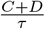 *bacycle* estimations (2-sided Wilcoxon signed-rank test on 36 pairs of values, p-value = 2.37 x10^−4^, effect size of 0.433). However, this statistically significant bias culminates at a difference in medians between paired samples of 0.0122, a value hardly significant from a biological and experimental point of view. Therefore, the lack of precision of 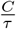 estimation does seem to bear no experimentally relevant impact on the precision of 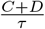 estimates on experimental data as well.

As an illustration, the effect of constraining or not 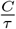 is shown in Figure 6C-E). The data are well represented by both models as shown by the overlap between the data distribution (±95% confidence interval) and the 95% credible interval for both models (Figure 6C). The cell cycle parameter space explored along the Nested Sampling optimization with loose (panel D) or tight (panel E) prior as a log-likelihood heatmap. Without constraining prior, 3 log-likelihood maxima (yellow regions) appear and 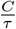 is improperly estimated by the median value of multiple mathematically optimal solutions (Figure 6D). When constraining 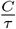 over the range of experimentally determined values (*µ*=1.09±0.1), a unique solution is found (Figure 6E). However, in both cases all potential solutions are associated with a very similar 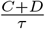 value.

#### Plasmids do not impair 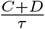 estimation with *bacycle*

One of the primary reason to develop *bacycle* is to allow cell cycle studies of natural (and clinical) bacterial isolates, or any bacterial strain for which the replication run-off treatment does not work effectively. One important feature of these strains regarding their DNA content is that most of them carry plasmids [49]. A back-of-the-envelope calculation suggests that plasmid DNA can represent as much as 5% of total cell DNA content, and could therefore alter cell cycle analysis.

We tested the impact of the presence of 2 well characterized plasmids, namely the pBR322 vector (4361bp, ∼20 copies/cell) and the F plasmid (∼100kb and 1 to 2 copies per cell). pBR322 has a relatively high copy-number for natural plasmids and F belongs to the most prevalent plasmid group found in *E. coli* strains [49]. We show here that the distribution of the DNA amount per cell with or without any combination of these 2 plasmids do not significantly change (Figure 7, B). In the absence of replication run-off, the estimation of 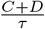 remains stable for all strains, regardless of the plasmid carried by the cells (Figure 7B) (Kruskal Wallis H-test on 4 groups, n=3 per group, p-value = 0.93).

**Fig 7.**
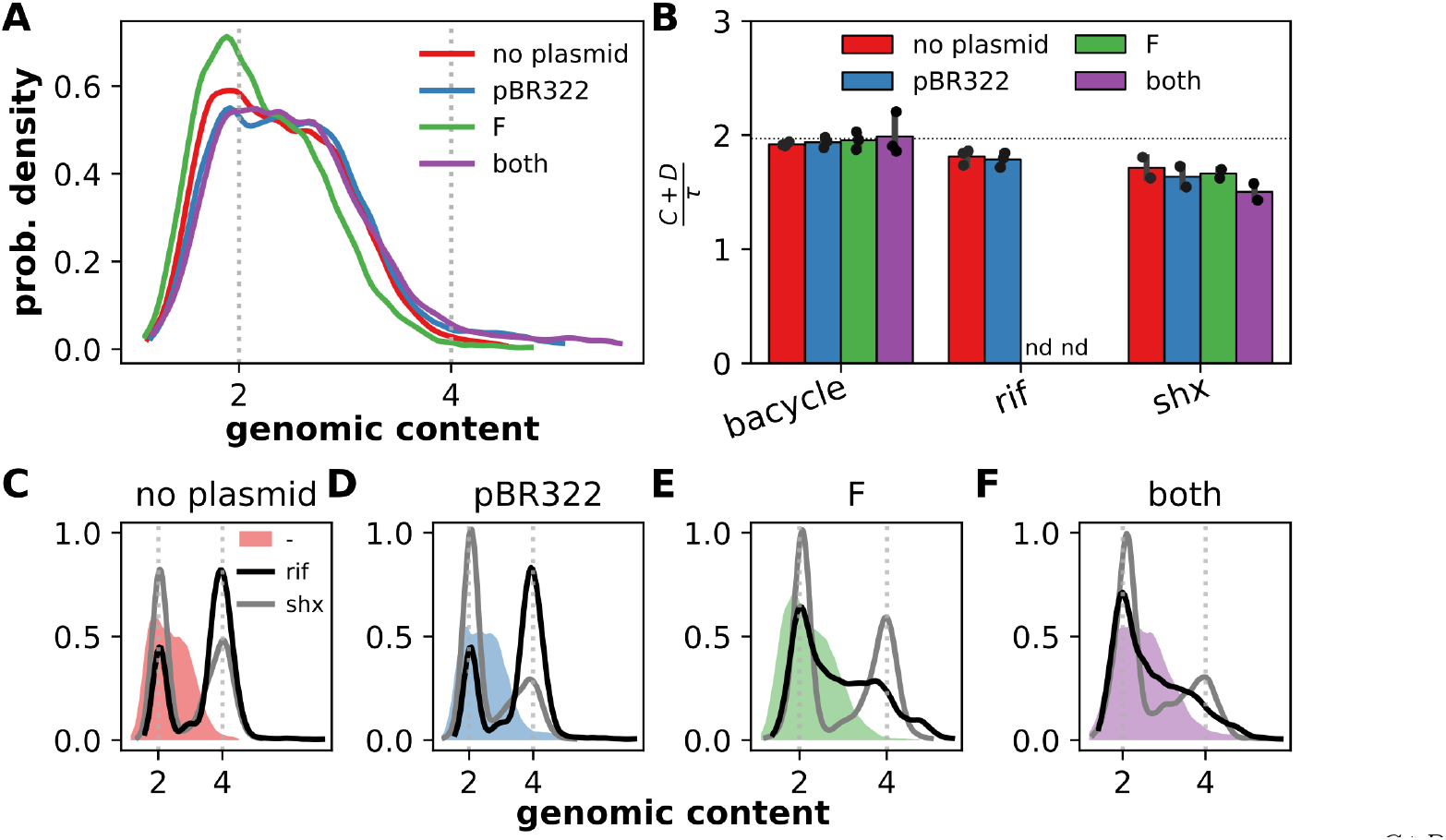
Effect of the presence of plasmids on DNA amount per cell and the estimation of 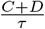. A) Distribution of DNA amount per cell quantified by flow cytometry for *E. coli* K12 MG1655 *dnaN::gfp-dnaN* carrying no plasmid, pBR322, F or both plasmids. Distributions were aligned according to the *f*_*c*_ parameter fitted with *bacycle*. B) Histogram showing the relative 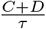 cell cycle period estimated with *bacycle* or using 2 different treatments to achieve replication run-off for the 4 plasmid conditions (color-coded). 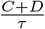 could not be estimated (nd: not determined) with the rifampicin-mediated replication run-off for cells carrying plasmid F alone or F and pBR322. Each replicate is indicated by a black dot (n=3, except for SHX treated samples: n=2). C-F) cytometry profiles before (filled curve) or after treatments (rifampicin: gray, SHX: black) for MG1655 *dnaN::gfp-dnaN* carrying no plasmid, pBR322, F, or both, respectively. The horizontal dashed black line indicate the average value obtained with *bacycle* for all 4 genetic conditions. Pharmacological treatments lasted for at least 3h. The color code reflect the plasmid carried by the cells in all panels of the figure. Distributions were aligned according to the *f*_*c*_ parameter fitted with *bacycle*.

When comparing the plasmid-free with pBR322 carrying cells, the estimated 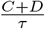 relative cell cycle period are in broad agreement, albeit estimations derived from replication run-off experiments tend to be smaller (2-way ANOVA on 16 values, Medium effect p-value = 0.52, Treatment effect p-value = 4.7110^−4^). *bacycle* estimates are greater than rifampicin mediated replication run-off 0.12 units, and than serine hydroxamate mediated replication run-off by 0.25 units (Figure 7). These differences represent 6.7% and 13% deviations,respectively, and remain rather small. Nonetheless, the differences between rifampicin and serine hydroxamate treatments reflect the different proportions of cells corresponding to 2 and 4 genome equivalents (Figure 7C, D). As reported earlier [14], both replication run-off methods are not equivalent.

Strikingly, the presence of the F plasmid seem to impede replication run-off and no clear separation of the 2 populations of cells with 2 or 4 genome equivalents could be achieved, whether the pBR322 plasmid was present or not, on top of the F plasmid (Figure 7E, F). However, even in the presence of the F plasmid, replication run-off could be achieved with serine hydroxamate + cephalexin in a similar manner, and with similar cytometry profiles (Figure 7C-F) and 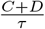 estimates (Figure 7B) as when the F plasmid is absent. The F plasmid effect is reminiscent of what has been observed in clinical *E. coli* isolates (Figure 1).

Despite the slight underestimation of 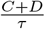 with rifampicin mediated replication run-off (6.7%), it seems that the presence of pBR322 or F plasmid do not strongly alter the cell cycle progression. In any case, the presence of these 2 plasmids does not seem to hamper cell cycle parameter estimation with *bacycle*.

## Discussion

The analysis of bacterial cell cycles is fundamental to understand bacterial growth and proliferation. We showed that state of the art methods for cell cycle analysis often fall short for clinical isolates of *E. coli* due to their highly heterogeneous response to rifampicin treatment. To address this issue, we propose a novel Python coded tool, *bacycle*, which proposes 2 complementary methods based on Monte Carlo approaches (Markov Chain Monte Carlo or Nested Sampling) to estimate cell cycle parameter values from cytometry profiles without drug treatment. These methods promise to enhance the accuracy, reliability and throughput of cell cycle analysis beyond laboratory strains and conditions.

State-of-the-art methods for cell cycle analysis in *E. coli* rely on replication run-off experiments which are based on the inhibitory effect of rifampicin on the initiation of DNA replication while allowing completion of ongoing replication cycles. We show here that this rifampicin effect is far from being common, even for bacterial strains of the same *E. coli* species. We could show that replication run-off experiments fail for 16 out of 20 clinical *E. coli* strains tested, distributed across 7 phylogroups (A, B1, B2, C, D, E and F, see Figure 1 and Table 1). The absence of narrow peaks in the distribution of DNA amount per cell at 2^n^ number of chromosome(s) (with n an integer value) suggests that the antibiotic treatment not only blocks the initiation of DNA replication but also impairs ongoing replication forks in some cases. We could show that the presence of F plasmid can induce the treatment failure in a strain that otherwise responds to the drug treatment (Figure 7C-F). Remarkably, the prevalence of F plasmids in strains of the *E. coli* species is high [49] and could explain the rifampicin-mediated replication run-off failure in a substantial fraction of *E. coli* strains, if we speculate that this plasmid effect is a general feature of the incompatibility plasmid group IncF. Nevertheless, some of the strains tested in this study did not respond to the rifampicin-mediated replication run-off treatment, despite the fact that they do not carry any IncF plasmid (or any plasmid at all, *e*.*g*., *E. coli* CFT073, see Figure 1B).

*bacycle* builds upon previous studies from Boye and collaborators, followed by Hansen and colleagues nearly 20 years later [18, 19], who showed that the Cooper-Helmstetter cell cycle model could be used to simulate distributions of DNA amount per cell. We therefore faced the same challenging limitation of the non-differentiability of this model. As a result, standard optimization procedures based on gradient descent are inefficient at identifying good parameter estimates. This limitation may have been the main reason why the models’ parameters were previously fitted manually. Here, we propose a Nested Sampling approach to overcome this limitation and prevent user-based differences in the appreciation of the goodness of the fit. Nested Sampling has also the advantage of readily pointing out situations where the model is not identifiable. This aspect of the optimization approach seems crucial in this case as a relatively small error on the *f*_*c*_ parameter (*>*20%) may lead to a large error in the biological parameter values (see Figure 3). This analysis of the identifiability of the model puts the emphasis on the definition of experimentally derived constrains for the fluorescence to DNA amount conversion factor (Figure 2).

We emphasize the importance of characterizing the limits of the tool even from a theoretical point of view. We investigated whether the model and the NS optimization algorithm implemented in the *bacycle* package did not introduce a lack of precision, or bias by design. When fitting parameters to data that were simulated with the underlying model, we showed that *bacycle* does not bias parameter values and provide estimates with a precision well below the experimental reproducibility (Figure 4 and S5A, B). In addition, we could document a not so uncommon limitation of models, that is the non-identifiability of the Cooper-Helmstetter model (Figure 3). Multiple sets of parameter values can describe the same cytometry profile by changing the scale parameter *f*_*c*_. Care should be taken to experimentally define a range of values for this fluorescence to DNA mount conversion factor so as to refine the parameter search within a parameter space region where the model can identify a unique solution (*i*.*e*., identifiable). We find that constraining the conversion factor *f*_*c*_ within ±20% of the experimentally determined value is usually enough to reach this identifiability goal. Care should be taken to verify that the solution returned by the algorithm is indeed unique by using the tools proposed in *bacycle*, or even more precisely, the ones included in the packages *dynesty* and *emcee* (*e*.*g*., corner plots).

The D, or C+D periods are experimentally difficult to measure and flow cytometry and replication run-off have been the state of the art technique to estimate them. The comparison of *bacycle* with replication run-off experiments shows that both approaches return highly similar results, both in terms of value and variability. First, the results obtained in this study with both methods are in agreement with previously published data collected for the *E. coli* K12 MG1655 strain over the last 20 years (see [19, 20, 50, 51], and Figure 5A). Second, the correlation between 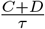 estimations with *bacycle* and replication run-off carried out on cells sampled from the same cultures is high (r=0.95, Figure 5B). Lastly, in our hands the reproducilibility of 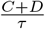 estimations falls below 10% of variations around the mean for both experimental approaches (Figure S5A).

We also analyzed cell cycle parameters with another strain expressing a fluorescent fusion with the *β*− clamp loader protein DnaN. With this strain we could show that the fluorescent fusion is not neutral to the progression of the cell cycle. As cells grow faster, replication run-off reveals some degree of asynchrony in the initiation of DNA replication that result in extra peaks in the distribution of DNA content per cell at unexpected numbers of genome equivalents (*3,5,6 or 7*, see Figure S5C-F). This degree of asynchrony slightly impairs the ability to estimate 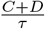 by replication run-off (Figure 5C). This effect adds up to the difficulty to analyze the distribution of fluorescence signal along the cell body as non-diffraction-limited GFP-DnaN foci appear (more replication forks) as cells grow faster. Thus, *bacycle* constitutes a unique tool providing quantitative estimation of the C+D period that is highly complementary to current methods and may overcome some of their limitations (requirement for drug-treatments or mutations to incorporate labeled nucleotides in the chromosome, effect of traductionnal fusions for fluorescence microscopy).

Concerning other parameters, we find that the estimation of the relative duration of the replicative period 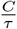 is less precise than the determination of 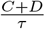. This lack of precision for the replication period is partly due to the fact that the fine structure of the distribution of the amount of DNA per cell, which is mainly due to the 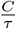 parameter (Figure 2A), can be captured and smoothed out with a higher *cv* value. The precision in 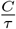 estimation can be greatly improved using drug-free experimental approaches, such as qPCR, digital PCR, or marker frequency analysis. We showed here how this complementary method can be used to constrain the prior on the 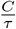 parameter in *bacycle* for fast growth conditions. *bacycle* allows for a good estimation of both biological parameters for slow growth conditions. Nevertheless, we could show the estimation of 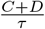 is achieved without loss or precision or increase in bias in the absence of this extra experimental knowledge (Figure S5A, B). This robustness of 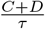 estimation is most likely due to the experimentally derived constrain on *f*_*c*_. We also noted an overestimation of the *cv* on experimental data, raising from 5 to 10% up to 20% from replication run-off samples to untreated samples. We speculate that this overestimation stems from 2 sources. As mentioned above, the fine structure of the distribution of DNA amount per cell in experimental cytometry profiles may be captured as noise and smoothed out through an increased *cv* value. We speculate that another probable source of noise could be the variability of the C and D periods in a population of cells that is not taken into account in the underlying model. This variability is bound to blur the ideal profile and mask the distribution fine structures and favor the increase of the *cv* value, at the expense of the 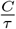 estimation.

A large fraction of bacterial isolates harbor extra-chromosomal replicons [52]. Here we show that 2 known plasmids – pBR322 and F – hardly affect the cell cycle dynamics and do not prevent the cell cycle analysis by flow cytometry (Figure 7A, B). However, the presence of the F plasmid, widely distributed in the *E. coli* species [49], seem to prevent the completion of the replication run-off (Figure 7C-F). Initiation of DNA replication at the origin (*oriC*) is indeed inhibited and that ongoing replication forks are maintained but cannot complete the chromosome replication as a relatively large fraction of cells with DNA contents in between the 2 characteristic peaks, at 2 and 4 genome equivalents under these growth conditions. We do not know how the presence of the F plasmid could alter the processivity or restart of replication forks in the presence of rifampicin, but not serine hydroxamate. It remains that beyond plasmids, it is estimated that nearly 15% of bacteria possess a multipartite genome with large replicons, called mega-plasmids or secondary chromosome, depending on the definition of these terms [52]. One important feature of these large secondary replicons is that their replication is integrated with the primary chromosome replication. As a general trend, these replicons are integrated within the cell life cycle by coupling the replication termination of the multiple replicons. So far, such replication patterns cannot be analyzed with *bacycle*, but could constitute an important future development to extend cell cycle studies to a wider range of clinically (*e*.*g*., *Vibrio cholerae*), biotechnologically (*e*.*g*., *Cupriavidus necator*) or agronomically (*e*.*g*., *Agrobacterium tumefaciens*) relevant organisms. The Cooper-Helmstetter model has been modified to identify a time delay between the initiation of replication of chromosomes I and II in *e*.*g*., *Vibrio cholerae* [53] and this type of modification could be used to objectively estimate this time delay from untreated cell samples using the Bayesian approaches

## Conclusion

We anticipate that the development of *bacycle* represents a significant advancement in the field of bacterial cell cycle analysis. By leveraging Nested Sampling and the Cooper-Helmstetter cell cycle model, *bacycle* overcomes the limitations of current methods based on replication run-off and provides accurate and reliable cell cycle parameters estimates from cytometry without any pharmacological treatment for clinical isolates of *E. coli*.

## Author Contributions

MC designed the research. AM, IB and MC carried out experiments. FS, MB and MC wrote the code and carried out simulations. AM, FS, ED and MC analyzed the data and wrote the article.

## Acknowledgments

We thank all the members of the [Genome Dynamics](https://cbi-toulouse.fr/equipe/bouet-cornet/) team and the [MAMBO](https://cbi-toulouse.fr/equipe/weiss/) team at the CBI for fruitful discussions. This work was supported by the grant ANR-20-CE13-0001-01 (MC).

## Supporting information

### *E. coli* strains are sensitive to rifampicin

**Fig S1.**
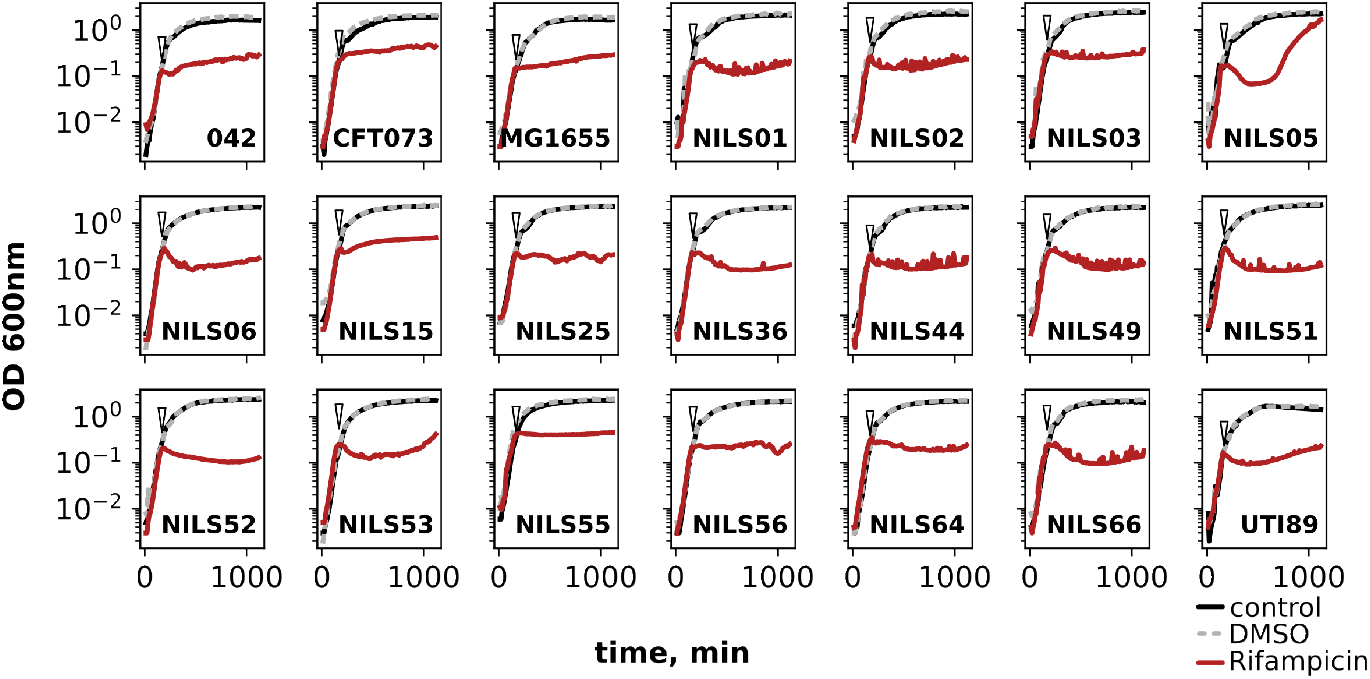
Growth arrests induced by rifampicin in *Escherichia coli* strains. Strain names are indicated in each panel. Black curves are reference growth curves without any antibiotic addition. Red lines show growth curves with the addition of rifampicin (300*µ*g/mL) at 3h after inoculation, as indicated by the black arrows. Dotted gray lines show growth cirves with the addicition of DMSO at the same time as rifampicin.

### Noise calibration with beads

Noise calibration with beads give similar results, albeit with different values. The fluorescence detection is not performed through the same channel et the same FRET procedure as for the DNA quantification procedure. Nonetheless, this approach verifies the value of ∼3% of the coefficient of variation for particle fluroescence detection, as stated by the constructor (3.6% - Figure S2 right panel)

### Non-identifiability of the Cooper-Helmstetter model

Non identifiability becomes even more relevant at higher biological parameter values, with more biologically relevant sets of parameters leading to identical cytometry profiles after scaling.

### Bias and precision from simulated cytometry profiles

Experimental and biological noises alter ideal measurements and estimation procedures must be robust to noise levels. But it is also important to verify that the estimation algorithm provides as well unbiased estimates, with high precision without noise, and then control how robust to noise are the bias and precision levels of the procedure. Data presented in Figure 4C are represented here over a grid of biological parameter values.

### Comparison of different experimental methods to estimate 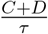

The drug-free approach developped in this study is compared to the state of the art replication run-off approach (see also Figure 5), as well as to microscopy based method using fluoresently labelled replication forks.

**Fig S2.**
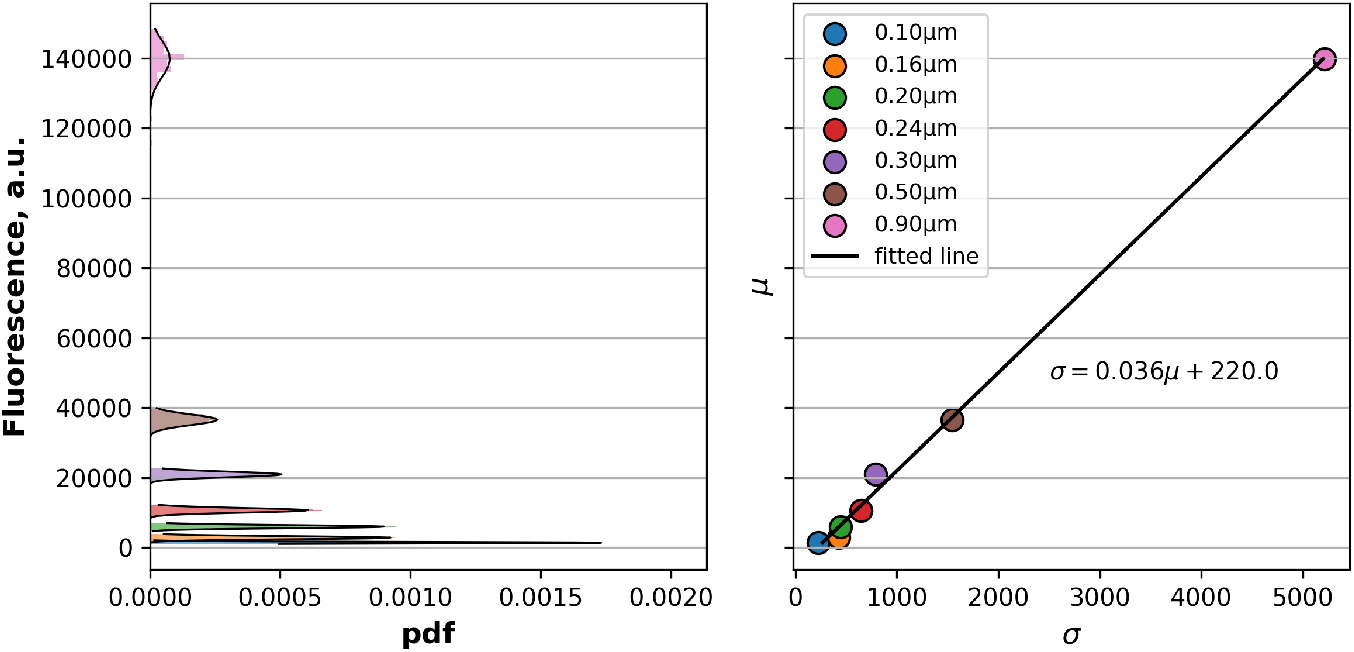
Noise calibration with fluorescent beads. Left panel: Probability density distribution of fluorescence associated beads of different sizes (color coded) at 520nm (emission wavelength). Normal distributions fitted to data are represented as black lines. Right panel: scatter plot of the standard deviation (*σ*) versus the mean (*µ*) of the Normal distributions fitted to the data represented on the left panel. The equation of the line fitted to the points indicates the respective contribution of Poisson (slope) and Gaussian (intercept) processes to the noise.

**Fig S3.**
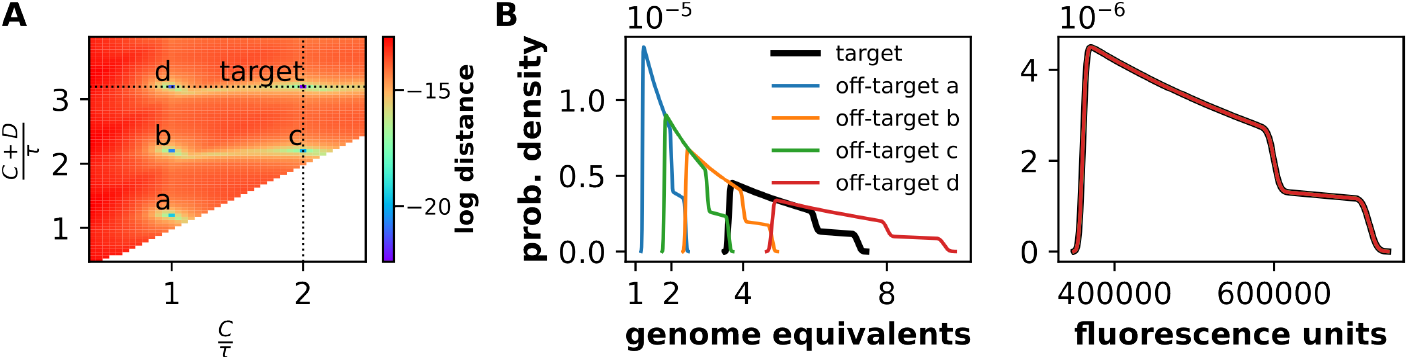
General non-identifiability of the Cooper-Helmstetter model. A) Matrix of distances between the target probability distribution function (pdf) simulated with parameters [2.0, 3.2, 0.01, 1.10^5^] with the pdf generated with other values for the first two parameters (biological parameters). The distance is colorcoded and is calculated as the Wasserstein distance and is independent of technical parameters. Three zones of short distances appear, indicated by the crossing of the 2 black dashed lines (true parameter values) and by the characters A and B (2 off-target solutions). Parameter space where 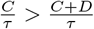 is not accessible by definition (white area). B) Illustration of the pdf corresponding to the 5 parameter sets in terms of cellular DNA contents on the left panel. The right panel shows that once shifted and normalized by the proper conversion factors, the distributions are indistinguishable. Conversion factor for off-target solutions reflect the changes in DNA amounts per cell induced by the deviation from the target values for both biological parameters by one or two units.

### Flow cytometry - gating strategy

#### Droplet digital PCR controls

The checklist of the Minimum Information for Publication of Quantitative Digital PCR Experiments is available online (DOI:10.5281/zenodo.18556582).

**Fig S4.**
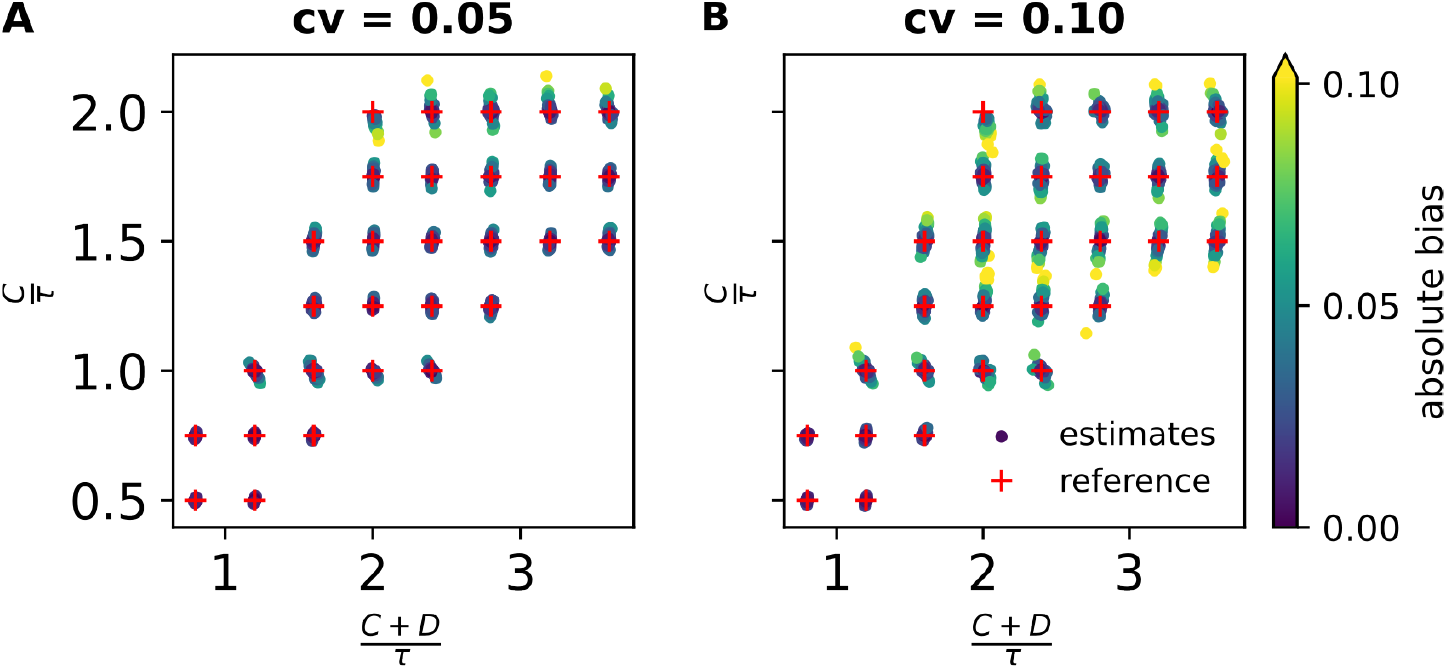
Bias and precision of Nested Sampling estimates from simulated, theoretical cytometry profiles. Each pair of parameter values distributed over the grid indicated by the thin dotted lines was independently tested 30 times and each dot represents one of these repeats. Marker colors illustrate the absolute bias on 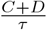, and were plotted such that repeats with the largest bias are plotted on top of those associated with smaller biases. Parameter space where 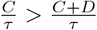 is not accessible by definition. The green area covers the range of possible values for *E. coli*.

**Fig S5.**
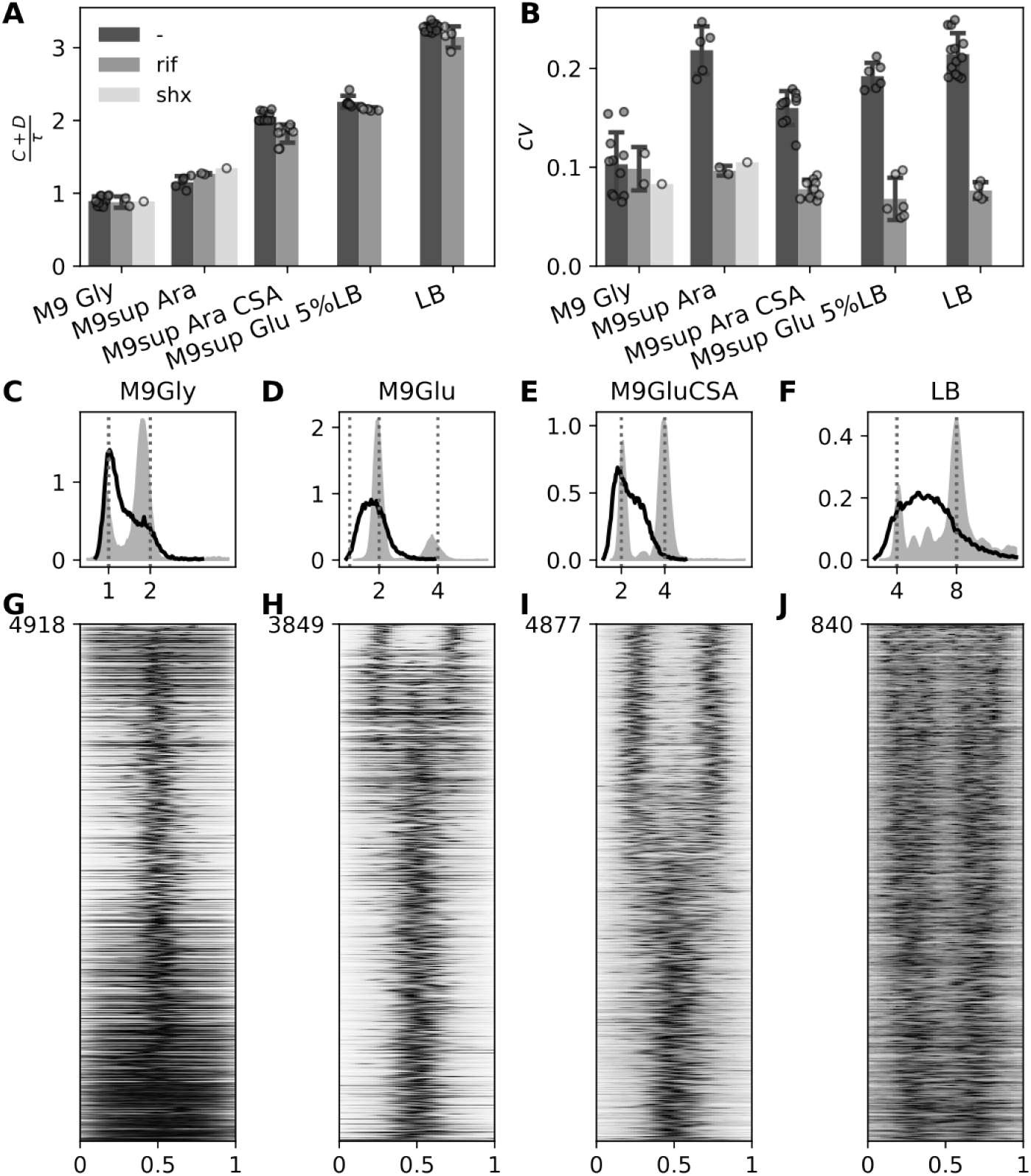
Cell cycle analysis with 3 methods on the same cell samples. A) Histogram and strip plot of all the 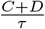 estimations with *bacycle*for the strain *E. coli* K12 MG1655 grown in different growth media. The grey shades correspond to the treatments (or absence thereof) A) Histogram and strip plot for the *cv* parameter estimated with *bacycle* for the same cytimetry profiles as in A). The color code is the same as in A). of all the 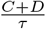 estimations with *bacycle*for the strain *E. coli* K12 MG1655 grown in different growth media. C-F) Distribution (probability density) of DNA amount per cell for the *E. coli K12* (*dnaN*)::*gfp-dnaN* under different growth conditions (growth media indicated as panel titles) and with (gray-filled curves) or without (black curves) replication run-off pharmacological treatment for the same cell samples (after or before treatment) for a representative experiment. The x-axis indicates the number of genome equivalents. G-J) Demographs associated with cytometry profiles illustrated in panels C to F, where each line represents the GFP-DnaN fluorescent signal from one cell pole to the other in relative units, with fluorescence profiles sorted from the shortest (bottom) to the longest (top) cell. Cell length is used as a proxy for cell age and the evolution of the GFP-DnaN fluorescence profile as a function of cell length provides an averaged view of the cell cycle from snap-shot images of an asynchronous population of cells. The fluorescence signal was normalized per cell so that the intensity each cell maximal intensity is set to 1 (black). The fluorescence intensity scales from 0 (white) to 1 (black). The number of cells analyzed for each demograph is indicated at the top left of each demograph.

**Fig S6.**
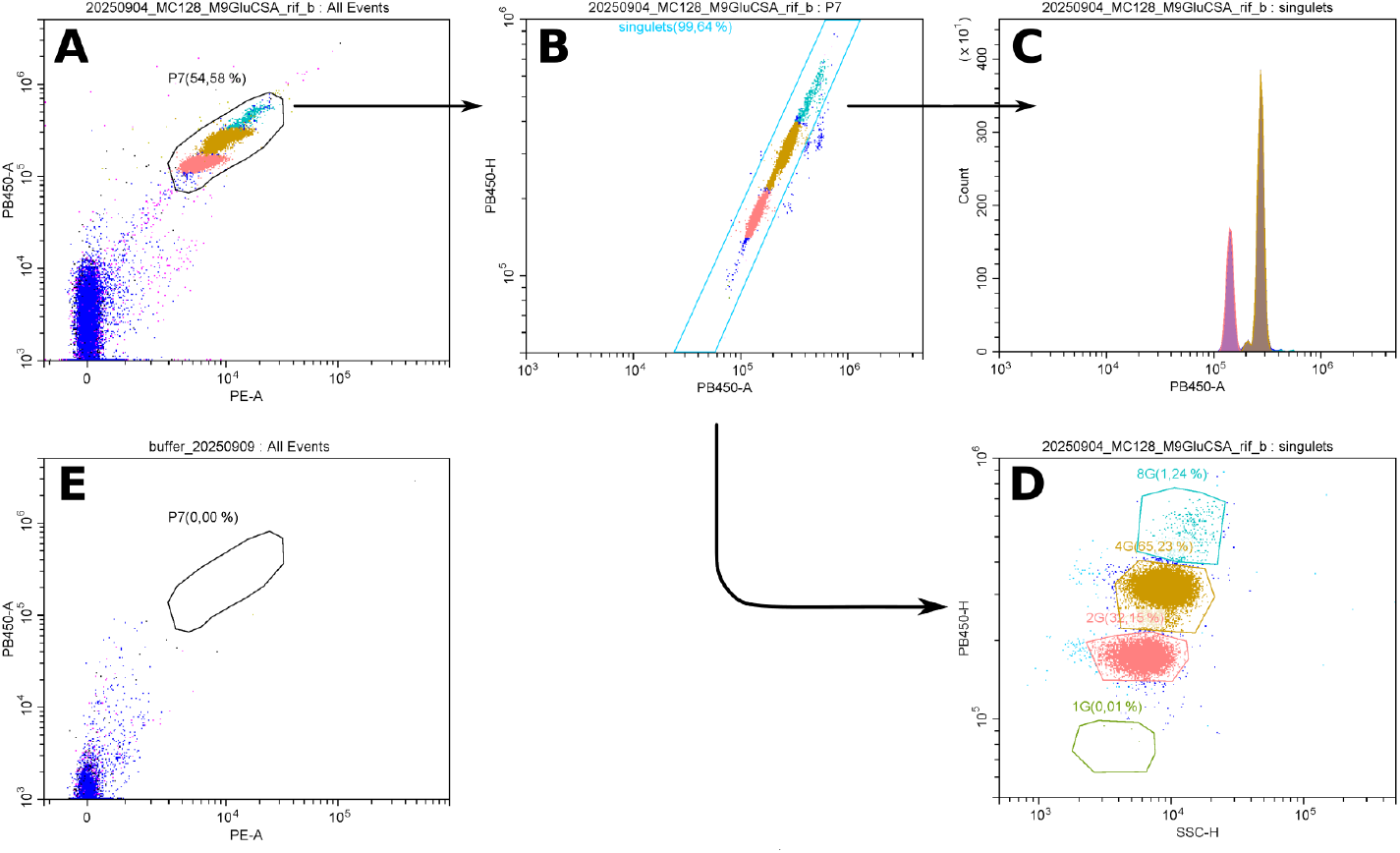
Gating strategy to extract particle of interest. A) All detected events are represented in a 2D plane representing the fluorescence amounts in the green (x-axis) and red (y-axis) channels. Particles with high fluorescence in both channels are considered as cells and a polygon is used to isolate cells (P7). B) Particles in P7 are represented in a red fluorescence signal Area (x-axis) versus Height (-y-axis). Particles out of the main diagonal, with higher Area signal than expected are considered as doublets and excluded and cells in the Singulet area are kept for further analysis. C) Gated particles in ‘singulets’ are used to construct DNA amount per cell distributions. D) The same particles can be further separated in the SSC-Height versus red fluorescence Height signal to gate particles with finite amount of genomes per cell (from 1 to 8 genome equivalents on this graph). E) The labeling buffer is analyzed for each cytometry session to verify that the buffer did not contain particles of the size of cells.

**Fig S7.**
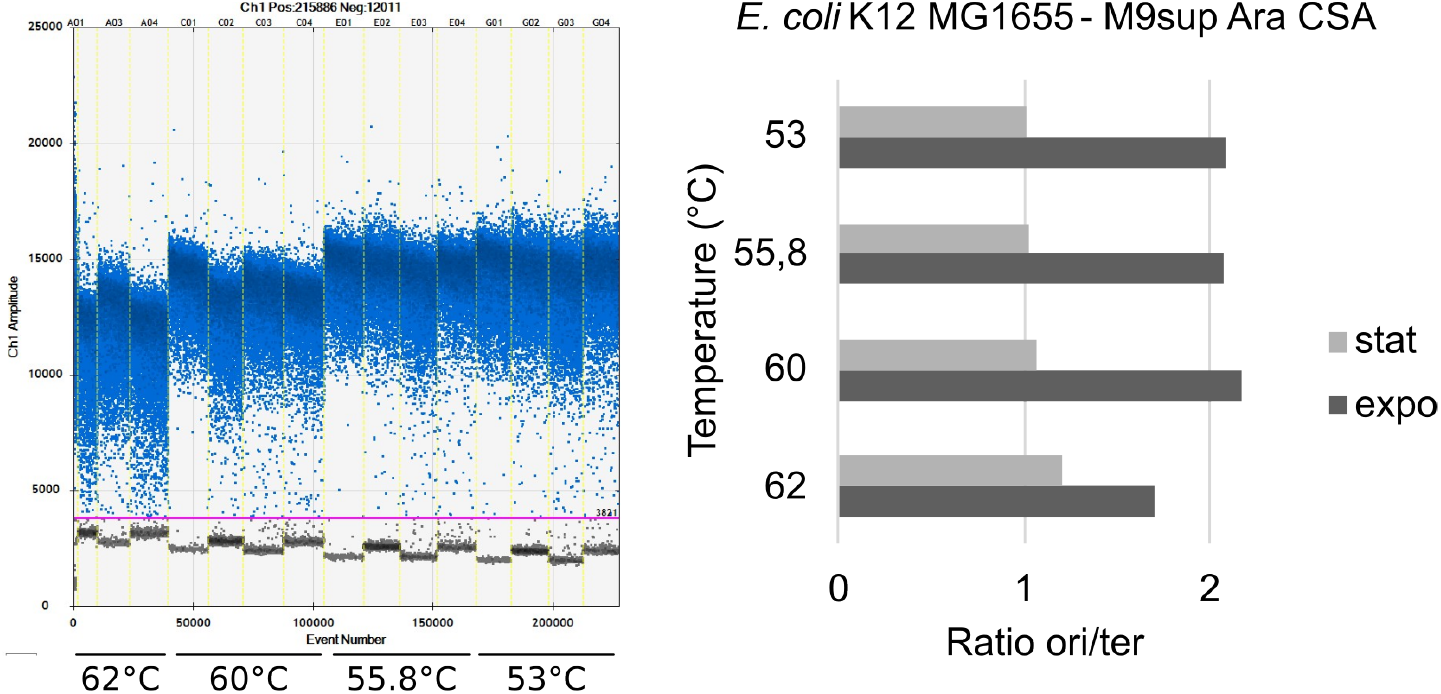
ddPCR thresholding and sensitivity to annealing temperature. The standard output of the QuantaSoft software (Biorad) provides a visual representation of the fluorescence level of each droplet (each dot) in the different reaction wells, separated by yeallow vertical dashed lines. The *ori* target is amplified in odd wells and the *ter* target in even wells. *E. coli* K12 MG1655 genomic DNA was purified from exponentially growing cells (columns 1 and 2) or stationnary phase cells (columns 3 and 4). Different rows correspond to different annealing tempratures. The temperature of the annealing step of the amplification program is indicated below the wells. The resulting *ori/ter* ratios are represented on the histogram on the right panel for the

